# *Drosophila* oocyte specification is maintained by the dynamic duo of microtubule polymerase Mini spindles/XMAP215 and dynein

**DOI:** 10.1101/2023.03.09.531953

**Authors:** Wen Lu, Margot Lakonishok, Vladimir I Gelfand

## Abstract

In many species, only one oocyte is specified among a group of interconnected germline sister cells. In *Drosophila melanogaster*, 16-cell interconnected cells form a germline cyst, where one cell differentiates into an oocyte, while the rest become nurse cells that supply the oocyte with mRNAs, proteins, and organelles through intercellular cytoplasmic bridges named ring canals via microtubule-based transport. In this study, we find that a microtubule polymerase Mini spindles (Msps), the *Drosophila* homolog of XMAP215, is essential for the oocyte fate determination. mRNA encoding Msps is concentrated in the oocyte by dynein-dependent transport along microtubules. Translated Msps stimulates microtubule polymerization in the oocyte, causing more microtubule plus ends to grow from the oocyte through the ring canals into nurse cells, further enhancing nurse cell-to-oocyte transport by dynein. Knockdown of *msps* blocks the oocyte growth and causes gradual loss of oocyte determinants. Thus, the Msps-dynein duo creates a positive feedback loop, enhancing dynein-dependent nurse cell-to-oocyte transport and transforming a small stochastic difference in microtubule polarity among sister cells into a clear oocyte fate determination.

**Significance statement:** Oocyte determination in *Drosophila melanogaster* provides a valuable model for studying cell fate specification. We describe the crucial role of the duo of microtubule polymerase Mini spindles (Msps) and cytoplasmic dynein in this process. We show that Msps is essential for oocyte fate determination. Msps concentration in the oocyte is achieved through dynein-dependent transport of *msps* mRNA along microtubules. Translated Msps stimulates microtubule polymerization in the oocyte, further enhancing nurse cell-to-oocyte transport by dynein. This creates a positive feedback loop that transforms a small stochastic difference in microtubule polarity among sister cells into a clear oocyte fate determination. Our findings provide important insights into the mechanisms of oocyte specification and have implications for understanding the development of multicellular organisms.

## Introduction

Mature oocytes, also known as eggs, are usually the largest cell of the whole body. To achieve a large cell size, in addition to *de novo* synthesis of new materials, oocytes acquire mRNAs, proteins, and organelles from the interconnected sister cells (1, 2). In many species, including the classic model organism *Drosophila melanogaster* as well as mammals, oocytes are specified among a group of interconnected cells, called germline cysts, after incomplete cytokinesis (3, 4). The rest of the sister cells transfer cytoplasmic materials to fast-growing oocytes before undergoing apoptosis (5, 6). Therefore, the oocyte represents a “winners take all” paradigm. The big standing question is how the oocyte maintains its “winner” position during development.

The ovary of *Drosophila* melanogaster provides a powerful system to address the question of oocyte fate maintenance due to the ample availability of genetic and cell biology tools. Within *Drosophila* ovaries, a germline stem cell divides to produce a cystoblast, which undergoes four rounds of cell divisions with incomplete cytokinesis to generate 16 interconnected sister cells (called cystocytes) connected by intercellular cytoplasmic bridges, the ring canals (Figure 1A) (7). Among the 16 interconnected cystocytes, one cell is specified as an oocyte, and the 15 sister cells undergo endoreplication, becoming polyploid nurse cells to “nurse” the oocyte (8). The oocyte is not randomly determined in the 16-cell cyst; instead, the oocyte is selected between the first two daughter cells (called pro-oocytes) after the first division of the cystoblast, with four ring canals in the center of the germline cyst (the dashed orange box, Figure 1A) (9). The oocyte and the 15 sister cells are then encapsulated by a layer of somatic epithelial cells, also known as follicle cells, to form an oval structure called the egg chamber (Figure 1A).

**Figure 1.**
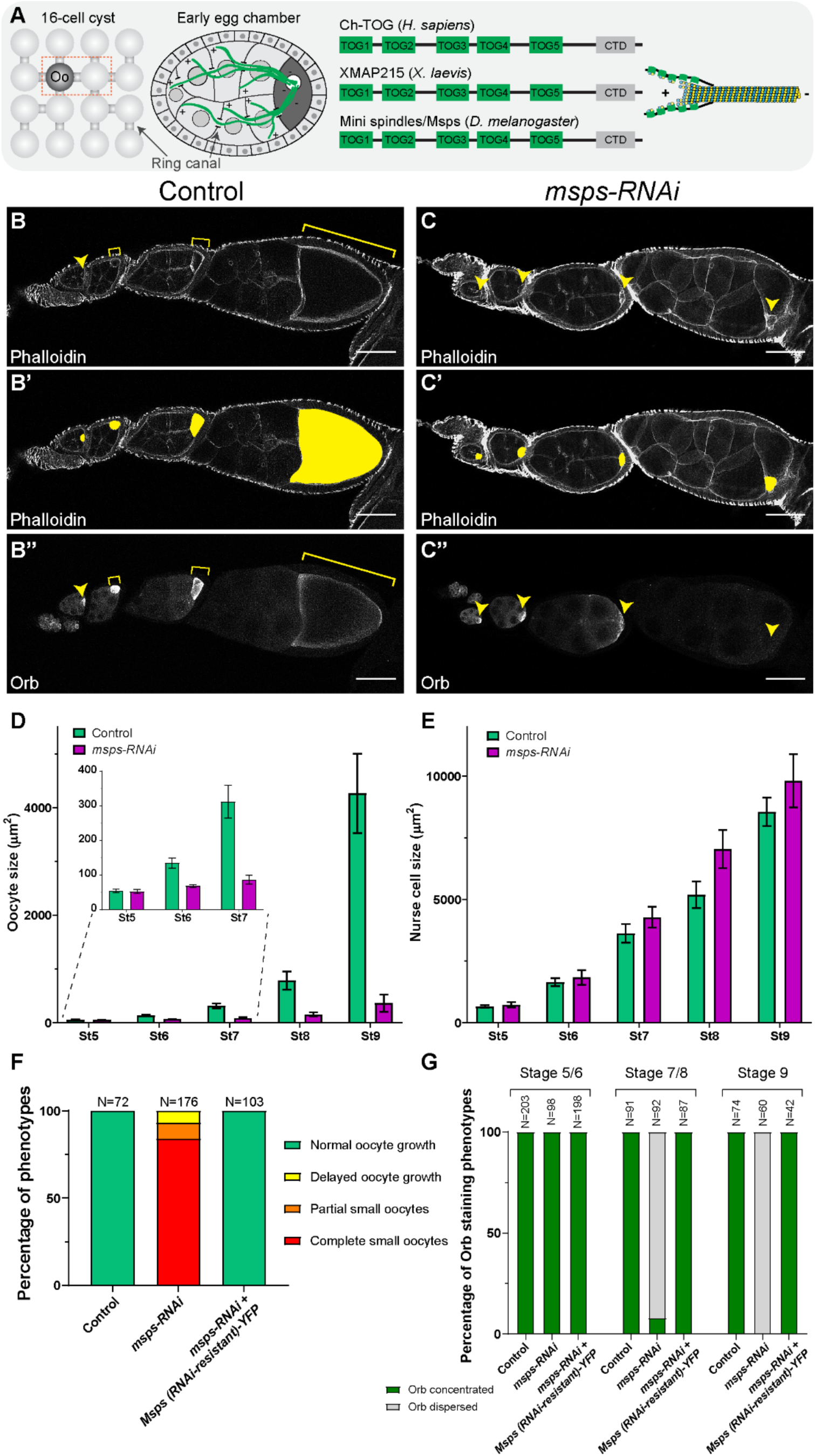
Msps is required for oocyte growth and fate maintenance. (A) A cartoon illustration of the oocyte specification in the germline cyst of 16-interconnected cystocytes. The oocyte candidates, the pro-oocytes, are outlined with the dashed orange box. Oo, oocyte. The microtubules in the early egg chamber display a highly polarized network, with minus-ends accumulated in the oocyte and the plus-ends extending into the interconnected nurse cells via the ring canals. The XMAP215-Dis1 family is a group of highly conserved microtubule-associated proteins that promote microtubule polymerization in eukaryotes. The homologies in *Drosophila* (Mini spindles, Msps), Xenopus (XMAP215), and human (Ch-TOG) all have five TOG domains that interact with either free tubulin dimer or microtubule lattice, with a C-terminal domain (CTD) is proposed to interact with End-binding protein EB1 to track the growing plus-ends of microtubules. (B-C’’) Phalloidin and Orb staining in control (B-B’’) and *Dhc64C-RNAi* (C-C’’) ovarioles. Oocytes are highlighted with the yellow arrowheads and brackets (B-C and B’’-C’’) or with yellow painting (B’-C’). Scale bars, 50 µm. (D) Measurement of oocyte sizes in stage 5 to stage 9 egg chambers. The *msps-RNAi* oocyte is significantly smaller than the control ones, starting stage 6. Control oocyte size: stage 5, 54.7 ± 5.6 µm^2^ (N=66); stage 6, 134.8 ± 14.8 µm^2^ (N=59); stage 7, 321.1 ± 47.6 µm^2^ (N=44); stage 8, 783.8 ± 166.0 µm^2^ (N=25); stage 9, 4263.8 ± 738.2 µm^2^ (N=20); *msps-RNAi* oocyte size: stage 5, 52.6 ± 5.3 µm^2^ (N=89); stage 6, 68.2 ± 4.6 µm^2^ (N=79); stage 7, 86.8 ± 12.8 µm^2^ (N=68); stage 8, 154.9 ± 37.2 µm^2^ (N=35); stage 9, 363.7 ± 159.7 µm^2^ (N=27). Data are represented as mean ± 95% confidence intervals. Unpaired t-tests with Welch’s correction were performed between control and *msps-RNAi* oocytes: stage 5, p= 0.5789 (not significant, n.s.); stage 6, p <0.0001 (****); stage 7, p <0.0001 (****); stage 8, p <0.0001 (****); stage 9, p <0.0001 (****). (E) Measurement of nurse cell sizes in stage 5 to stage 9 egg chambers. In contrast to the drastic reduction in the oocyte size, the nurse cells in *msps-RNAi* are not smaller than the control ones; instead, *msps-RNAi* nurse cells at stages 7-9 are larger than the control. Control nurse cell size: stage 5, 658.6 ± 60.2 µm^2^ (mean ± 95% confidence intervals) (N=66); stage 6, 1642.5 ± 158.9 µm^2^ (N=59); stage 7, 3632.6 ± 377.2 µm^2^ (N=44); stage 8, 5184.8 ± 538.2.0 µm^2^ (N=25); stage 9, 8559.7 ± 577.7 µm^2^ (N=20); *msps-RNAi* nurse cell size: stage 5, 730.8 ± 100.6 µm^2^ (N=89); stage 6, 1837.5 ± 303.3 µm^2^ (N=79); stage 7, 4277.5 ± 425.6 µm^2^ (N=68); stage 8, 7044.8 ± 425.6 µm^2^ (N=35); stage 9, 9823.1 ± 1080.1 µm^2^ (N=27). Data are represented as mean ± 95% confidence intervals. Unpaired t-tests with Welch’s correction were performed between control and *msps-RNAi* nurse cells: stage 5, p= 0.2223 (not significant, n.s.); stage 6, p= 0.2588 (n.s.); stage 7, p = 0.0249 (*); stage 8, p= 0.0002 (***); stage 9, p = 0.0398 (*). (F) Percentages of oocyte growth phenotypes in the listed genetic background (all with one copy of *maternal αtub-Gal4^[V37]^*). Classifications of oocyte growth phenotypes were previously described (27). For control samples: normal oocyte growth, 100%; delayed oocyte growth, 0%; partial small oocytes, 0%; complete small oocytes, 0% (N=72). For *msps-RNAi* samples: normal oocyte growth, 0%; delayed oocyte growth, 6.8%; partial small oocytes, 9.1%; complete small oocytes, 84.1% (N=176). For *msps-RNAi* + *Msps (RNAi-resistant)-YFP* samples: normal oocyte growth, 100%; delayed oocyte growth, 0%; partial small oocytes, 0%; complete small oocytes, 0% (N=103). (G) Percentages of the Orb staining phenotypes in stage 5-6, stage 7-8, and stage 9 egg chambers in listed genotypes (all with one copy of *maternal αtub-Gal4^[V37]^*). Characterizations of Orb concentration and Orb dispersion were previously described (27). For control samples: stage 5-6, 100% Orb concentrated and 0% Orb dispersed (N=203); stage 7-8, 100% Orb concentrated and 0% Orb dispersed (N=91); stage 9, 100% Orb concentrated and 0% Orb dispersed (N=74). For *msps-RNAi* samples: stage 5-6, 100% Orb concentrated and 0% Orb dispersed (N=98); stage 7-8, 7.6% Orb concentrated and 92.4% Orb dispersed (N=92); stage 9, 0% Orb concentrated and 100% Orb dispersed (N=60). For *msps-RNAi* + *Msps (RNAi-resistant)-YFP* samples: stage 5-6, 100% Orb concentrated and 0% Orb dispersed (N=198); stage 7-8, 100% Orb concentrated and 0% Orb dispersed (N=87); stage 9, 100% Orb concentrated and 0% Orb dispersed (N=42).

Microtubules and the associated proteins play a central role in oocyte differentiation. The formation of a single microtubule organization center (MTOC) within the 16-cell cyst is a hallmark of oocyte specification (10, 11). Depolymerization of microtubules by drugs such as colchicine results in the formation of 16 nurse cells and no oocyte in the cyst (10, 12). The centrosomes from nurse cells and the microtubule mins-end binding protein Patronin are both known to be accumulated in early oocytes (13, 14). The microtubules, in turn, are nucleated from the MTOC in the oocyte and extend their plus-ends into the interconnected nurse cells via the ring canals (Figure 1A) (15, 16). This polarized microtubule network is proposed to be essential for accumulating oocyte-specific factors by the action of the major microtubule minus-end-directed motor, cytoplasmic dynein (15, 16).

The components of the cytoplasmic dynein complex (hereafter referred to as “dynein”), including dynein heavy chain (Dhc64C), dynein light intermediate chain (Dlic), dynein light chain (Dlc), its key cofactors dynactin and Lissencephaly-1 (Lis1), its activator Bicaudal D (BicD), and its mRNA-binding adaptor Egalitarian (Egl) are all concentrated in the oocyte and are essential for the oocyte specification. Early genetic disruptions of Dhc64C, Dlc, Dlic, dynactin p150/Glued and p50/Dmn, Lis1, BicD, and Egl lead to the lack of oocyte specification in the 16-cell cyst (17–26). Dynein is not only required for oocyte specification but is also indispensable for maintaining oocyte differentiation. Later or weaker inhibition of dynein or its cofactors (such as Dhc64C, Dlic, p150/Glued, Lis1, Dlc, and Egl) results in gradual loss of oocyte identity (21, 27), indicating that maintenance of the oocyte differentiation requires continuous dynein-driven activity.

Intriguingly, dynein is not just working as a minus-end-directed motor along the pre-formed polarized microtubule network originating from the MTOC in the oocyte; rather, it is involved in organizing and maintaining this polarized network itself. The formation of MTOC is either blocked or disrupted in *BicD* and *egl* mutants (10, 13, 28). BicD is an adaptor that activates the motility of the dynein-dynactin complex and links the complex to its cargoes (29, 30), whereas Egl, an RNA binding protein, serves as a key adaptor for dynein-dependent transport and is essential for the localization of the polarity determinants (e.g., *oskar*, *gurken*, and *bicoid* mRNAs) in oocytes and early embryos (31–36). It raises a possibility that dynein transports some mRNA to the oocyte via its interaction with Egl, which in turn increases microtubule polymerization in the oocyte and promotes the oocyte fate determination and maintenance.

Microtubules are polymers composed of αβ tubulin dimers. Microtubules can dynamically undergo growing or shrinking. The growth of microtubules can be enhanced by a group of proteins, XMAP215/Dis1 family, that are highly conserved in eukaryotes. The XMAP215 (*Xenopus* microtubule-associated protein 215) was first identified in frog oocytes as a processive microtubule polymerase that stimulates microtubule growth *in vitro* (37, 38). The higher eukaryotic XMAP215/Dis1 family members, including the *Xenopus* XMAP215, its *Drosophila* homolog Mini spindles (Msps), and the human homolog chTOG, all possess an N-terminal array of five TOG (tumor <u>o</u>verexpressed <u>g</u>ene) domains (Figure 1A). TOG domain is composed of six HEAT repeats and forms a flat, paddle-like structure, and the intra-HEAT repeat loops are important for tubulin binding (39–41). The five TOG domains have evolved to have differential preferences for binding either free or polymerized tubulin: TOG1-3 bind to free tubulin dimers, while TOG4-5 prefer tubulin incorporated into the microtubule lattice (42, 43). XMAP215/Msps tracks the plus-ends of growing microtubules (42) via the interaction with the EB1 binding proteins SLAIN2 (44) and Sentin (45). Altogether, XMAP215/Msps uses a polarized array of TOG domains at the microtubule plus-ends and facilitates the addition of soluble tubulin dimers to the microtubule polymer, in which TOG1-2 interact with free tubulin and TOG5 binds to the microtubule lattice-incorporated tubulin, while TOG3-4 bridge and stabilize the intermediate conformation between free and incorporated tubulins (Figure 1A) (42). Knockdown of XMAP215/Msps in *Drosophila* results in shorter or disorganized spindles in mitotic and meiotic cells (46–48). In hypomorphic mutants of *XMAP215/Msps*, the defects in microtubule organization and mRNA localization during mid-oogenesis have been reported (49). However, due to its requirement in spindle formation and mitotic progression, the role of Msps in early oocyte development remained unclear.

Here we report that the *Drosophila* microtubule polymerase XMAP215/Msps is essential for oocyte determination and growth. Dynein transports and concentrates the *msps* mRNA in the oocyte. As a result, translated Msps protein accumulates in the oocyte and is retained via its interaction with oocyte microtubules. Msps promotes microtubule polymerization in the oocyte as well as microtubule growth from the minus-ends in the oocyte into nurse cells. The oocyte concentration of Msps increases the number of correctly-oriented microtubule tracks in the ring canals and thus increases dynein-dependent transport from nurse cells to the oocyte. Therefore, Msps and dynein together form a positive feedback loop that ensures more microtubules originate in the oocytes, and dynein transports more oocyte-specific mRNAs, proteins, and organelles into the oocyte. We propose that the future oocyte is the pro-oocyte stochastically with slightly more microtubules minus-ends, and it relies on the Msps-dynein duo team to amplify the microtubule polarity difference and thus maintain the oocyte differentiation.

## Results

### XMAP215/Msps is essential for oocyte growth and fate maintenance

The high microtubule polymerization in the oocyte has been well documented both in early (14) and mid-oogenesis (50). Here we aim to understand how this high microtubule polymerization activity is achieved. We tested the role of a microtubule polymerase, XMAP215/Msps, in the *Drosophila* germ line. As XMAP215/Msps is required for proper mitotic spindle formation, knockdown of Msps by an RNAi line under an early germline driver, *nanos-Gal4^[VP16]^* that is expressed in the primordial germ cells (51), results in the complete germless ovary (Supplementary Figure 1A-1C). To bypass the requirement of Msps in cell division, we employed a postmitotic germline-specific driver, *maternal α-tubulin-Gal4^[V37]^*, that starts the expression in egg chambers at stages 3-4, after the completion of cystocyte cell division (27, 31, 50). The postmitotic knockdown of XMAP215/Msps caused a complete oocyte growth arrest in most ovarioles (Figure 1B-1F; Supplementary Figure 1D). The oocyte remains small over the stages and fails to acquire cytoplasmic contents from the interconnected nurse cells (hereafter referred to as the ‘small oocyte’ phenotype). Interestingly, the oocyte marker, Orb (oo18 RNA-binding protein) (52) is properly concentrated in the oocyte during early oogenesis but gets completely dispersed in mid-oogenesis (Figure 1C-1C’’ and 1G). It indicates that *msps-RNAi* driven by *maternal α-tubulin-Gal4^[V37]^* does not interfere with the early oocyte specification but affects the oocyte fate maintenance.

To ensure that the small oocyte and oocyte cell fate loss are specific to the *msps* knockdown, we generated a YFP-labeled full-length Msps that carries silent mutations making it resistant to the RNAi line we used. This RNAi-resistant Msps-YFP was able to fully rescue the small oocyte and oocyte fate loss phenotypes caused by *msps-RNAi* (Figure 1F-1G; Supplementary Figure 1E-1E’), indicating that the oocyte growth and maintenance defects are caused specifically by lack of Msps, rather than off-target effects of *msps-RNAi*.

### XMAP215/Msps promotes microtubule polymerization in the oocyte

Msps belongs to the XMAP215 family and has been shown to promote microtubule polymerization in *Drosophila* cells (42, 43). Having confirmed that Msps is required for oocyte growth and fate maintenance, we proceed to examine the effect of *msps-RNAi* on microtubules in the germ line. We used live imaging of EB1, the end-binding protein 1 that tracks the polymerizing microtubule plus-ends (53), as a readout of microtubule polymerization. An EB1-GFP line under a *ubiquitin* promoter (*ubi-EB1-GFP*) (50, 54–56) showed that in wild-type egg chambers, the microtubule polymerization activity is much higher in the oocyte than in the interconnected sister nurse cells (Figure 2A and 2C-2D) (Video 1). This high microtubule polymerization activity results in more microtubule plus-ends growing from the oocyte into nurse cells (Video 2), which is important for establishing the polarized microtubule network for nurse-to-oocyte transport (Figure 1A). Knockdown of *msps* by RNAi leads to a drastic reduction of EB1 tracks in the oocytes and nurse cells, as well as in the nurse cell-to-oocyte ring canals (Figure 2B-2D) (Video 3). As a result, the amount of microtubules in the *msps-RNAi* oocyte is severely reduced compared to the control oocyte, as seen in both total tubulin antibody staining and the *in vivo* labeling with GFP-tagged Ensconsin/MAP7 microtubule-binding domain (EMTB-3XGFP) (27) (Figure 2E-2G; Supplementary Figure 2A-2C). Very intriguingly, the microtubule level in nurse cells is largely unaffected (Figure 2H; Supplementary Figure 2D), suggesting that most microtubules in nurse cells are stable and do not rely on microtubule dynamics. Together, these data show that Msps is the main factor driving high microtubule polymerization activity in the oocyte but not in nurse cells.

**Figure 2.**
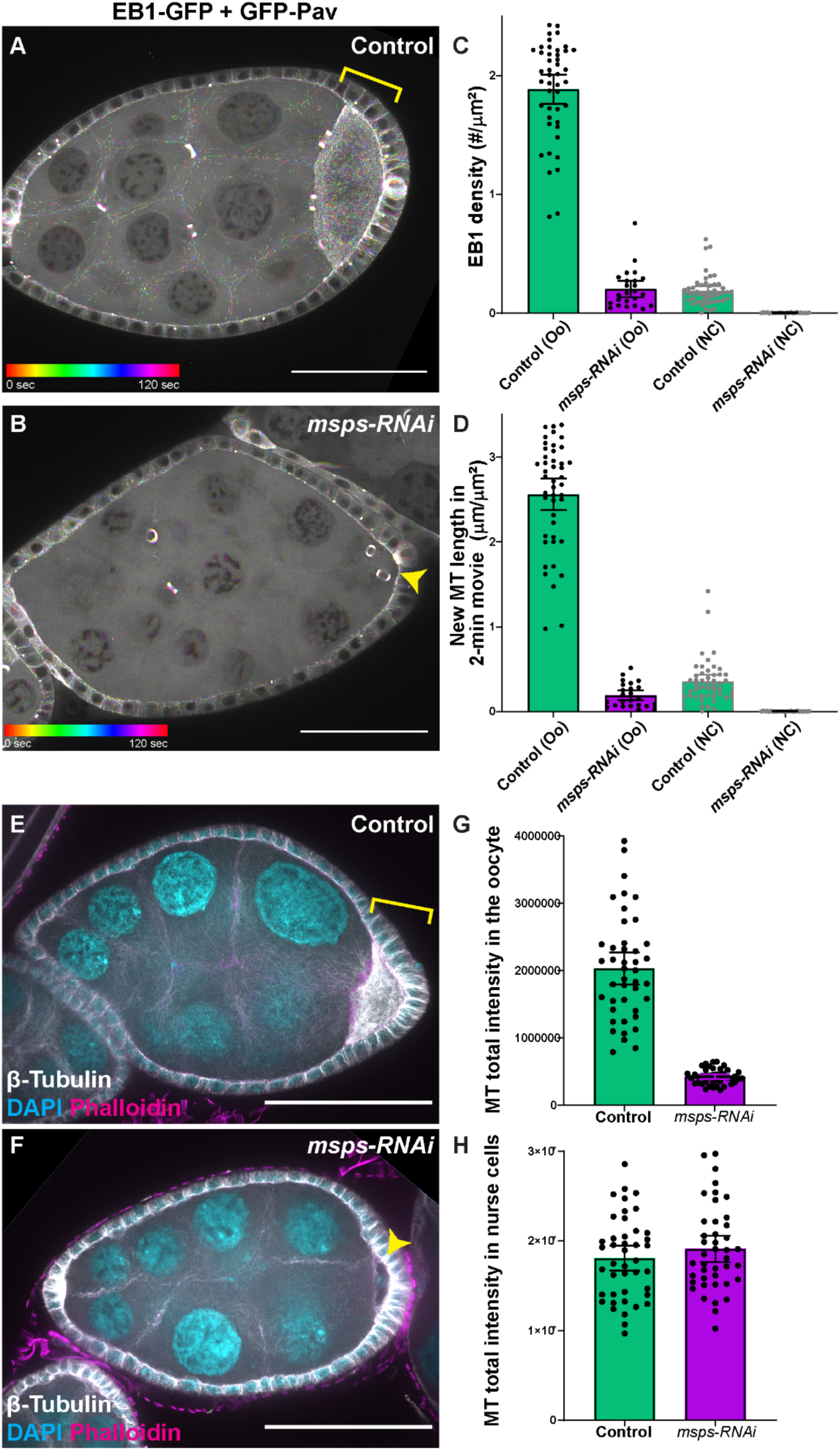
Msps promotes microtubule polymerization in the oocyte. (A-B) Temporal color-coded hyperstacks of *ubi*-EB1-GFP in control (A) and *msps-RNAi* (B). In control, more polymerizing microtubule plus-ends were seen in the oocyte than in nurse cells, indicated by the colorful tracks (A). In *msps-RNAi*, very few EB1 comets can be seen in the oocyte and nurse cells (B). In addition to *ubi*-EB1-GFP, a GFP-tagged Pavarotti/kinesin-6 under an *ubi* promoter (*ubi*-GFP-Pav) was used to illustrate the positions of ring canals. (C-D) Quantification of the density of EB1 comets and the length of newly polymerized microtubules in stage 6-7 egg chambers of control and *msps-RNAi*. Oo, oocyte, black dots; NC, nurse cells, grey dots. (C) The EB1 density in following samples: control oocyte, 1.887 ± 0.122 (/µm^2^) (N=45); *msps-RNAi* oocyte, 0.205 ± 0.069 (/µm^2^) (N=24); control nurse cells, 0.190 ± 0.038 (/µm^2^) (N=47); *msps-RNAi* nurse cells, 0.004 ± 0.001 (/µm^2^) (N=25). Unpaired t-tests with Welch’s correction were performed: between control and *msps-RNAi* oocytes, p<0.0001 (****); between control and *msps-RNAi* nurse cells, p<0.0001 (****). (D) The length of new microtubules (within 2 min): control oocyte, 2.562 ± 0.188 (µm /µm^2^) (N=45); *msps-RNAi* oocyte, 0.193 ± 0.060 (µm /µm^2^) (N=24); control nurse cells, 0.356 ± 0.077 (µm /µm^2^) (N=47); *msps-RNAi* nurse cells, 0.004 ± 0.001 (µm /µm^2^) (N=25). Unpaired t-tests with Welch’s correction were performed: between control and *msps-RNAi* oocytes, p<0.0001 (****); between control and *msps-RNAi* nurse cells, p<0.0001 (****). (E-H) Representative images (E-F) and quantification (G-H) of microtubule staining in control and *msps-RNAi* egg chambers. Total microtubule intensity in *msps-RNAi* oocytes is heavily diminished compared to control oocytes, while nurse cell microtubules are not largely affected by the *msps* knockdown. (G) Total microtubule intensity in oocytes: control stage 6-7, 2.03 ± 0.24 x10^6^ A.U. (arbitrary unit) (N=43); *msps-RNAi* stage 6-7, 4.2 ± 0.3 x10^5^ A.U. (N=42). Unpaired t-tests with Welch’s correction were performed between control and *msps-RNAi* oocytes: p <0.0001 (****). (H) Total microtubule intensity in nurse cells: in control stage 6-7 oocyte, 1.81 ± 0.14 X10^7^ A.U. (N=43); in *msps-RNAi* stage 6-7 oocyte, 1.91 ± 0.15 X10^7^ A.U. (N=42). Unpaired t-tests with Welch’s correction were performed between control and *msps-RNAi* nurse cells: p = 0.3042 (n.s.). Oocytes are indicated by the yellow arrowheads or yellow brackets. Scale bars, 50 µm. Data are represented as scattered individual data points with mean ± 95% confidence intervals (C-D, G-H).

Furthermore, we overexpressed a GFP-tagged Klp10A, the *Drosophila* kinesin-13 homolog known to depolymerize microtubules in *Drosophila* cells (57–59). Overexpression of Klp10A driven by *maternal αtub-Gal4^[V37]^* leads to significant microtubule loss both in the oocyte and nurse cells and results in the majority of the ovarioles having small oocytes and gradually losing oocyte fate, phenocopying the *msps-RNAi* mutant (Supplementary Figure 2E-2I). It suggested that the loss of microtubules is the main reason attributed to the defects in oocyte growth and fate maintenance observed in *msps-RNAi*.

### *XMAP215/msps* mRNA is concentrated in the oocyte by dynein-dependent transport

How does Msps differentially regulate microtubule polymerizations in the oocyte and nurse cells? To address this question, we decided to visualize *msps* mRNA using single molecule inexpensive fluorescence *in situ* hybridization (smiFISH) and found that *msps* mRNA is highly concentrated in the oocytes (Figure 3A-3A’ and 3D-3E). The smiFISH signal is specific to *msps* mRNA as germline knockdown of *msps* by RNAi leads to a complete abolishment of the smiFISH signal of *msps* RNA (Supplementary Figure 3A-3B’).

**Figure 3.**
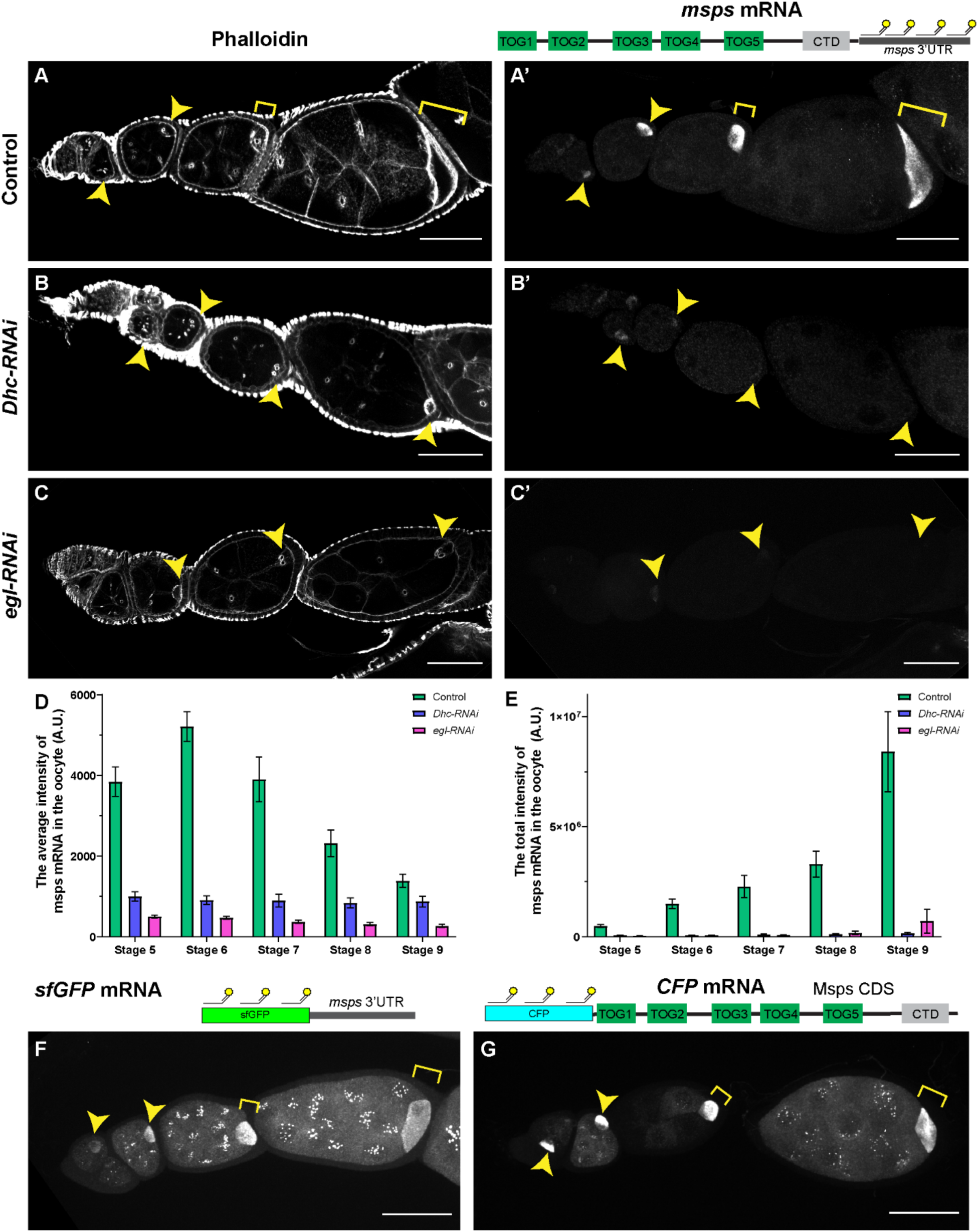
Dynein is required for *msps* mRNA concentration in the oocyte. (A-C’) Phalloidin and smiFISH staining of msps mRNA in control (A-A’), *Dhc-RNAi* (B-B’), and *egl-RNAi* (C-C’). Cy5-labeled FLAP-X smiFISH probes recognize the 3’UTR of *msps* mRNA. *msps* mRNA is concentrated in control oocytes but heavily reduced in *Dhc-RNAi* and *egl-RNAi* oocytes. (D) Average fluorescence intensity of *msps* mRNA (by smiFISH against *msps* 3’UTR) in control, *Dhc-RNAi*, and *egl-RNAi*. The average *msps* mRNA intensity in control oocytes: stage 5, 3842.6 ± 365.1 A.U. (N=38); stage 6, 5208.2 ± 367.1 A.U. (N=41); stage 7, 3903.1 ± 552.1 A.U. (N=19); stage 8, 2317.8 ± 325.8 A.U. (N=23); stage 9, 1390.5 ± 161.7 A.U. (N=24). The average *msps* mRNA intensity in *Dhc-RNAi* oocytes: stage 5, 996.5 ± 113.1 A.U. (N=54); stage 6, 908.6 ± 106.7 A.U. (N=47); stage 7, 896.2 ± 158.3 A.U. (N=38); stage 8, 837.0 ± 122.4 A.U. (N=32); stage 9, 871.2 ± 135.0 A.U. (N=30). The average *msps mRNA* intensity in *egl-RNAi* oocytes: stage 5, 500.8 ± 35.0 A.U. (N=71); stage 6, 469.4 ± 34.5 A.U. (N=56); stage 7, 369.8 ± 39.0 A.U. (N=55); stage 8, 308.3 ± 37.8 A.U. (N=32); stage 9, 270.4 ± 43.0 A.U. (N=25). Unpaired t-tests with Welch’s correction were performed between the following groups: control and *Dhc-RNAi* stage 5-stage 9, all p<0.0001 (****); control and *egl-RNAi*, stage 5-stage 9, all p<0.0001 (****); *Dhc-RNAi* and *egl-RNAi*, stage 5-stage 9, all p<0.0001 (****). (E) Total fluorescence intensity of *msps* mRNA (by smiFISH against *msps* 3’UTR) in control, *Dhc-RNAi*, and *egl-RNAi*. The total *msps* mRNA intensity in control oocytes: stage 5, 482091.0 ± 71442.2 A.U. (N=38); stage 6, 1494695.9 ± 211967.4 A.U. (N=41); stage 7, 2282849.2 ± 504102.6 A.U. (N=19); stage 8, 3292804.4 ± 590320.8 A.U. (N=23); stage 9, 8410028.3 ± 1821593.5 A.U. (N=24). The total *msps* mRNA intensity in *Dhc-RNAi* oocytes: stage 5, 65031.2 ± 7828.3 A.U. (N=54); stage 6, 65176.3 ± 7707.7 A.U. (N=47); stage 7, 94359.4 ± 21229.0 A.U. (N=38); stage 8, 110487.8 ± 24462.6 A.U. (N=32); stage 9, 146230.2 ± 52093.7 A.U. (N=30). The total *msps mRNA* intensity in *egl-RNAi* oocytes: stage 5, 43861.3 ± 3831.3 A.U. (N=71); stage 6, 55754.7 ± 4478.7 A.U. (N=56); stage 7, 78525.1 ± 10578.5 A.U. (N=55); stage 8, 166950.2 ± 87307.5 A.U. (N=32); stage 9, 703163.2 ± 543799.2 A.U. (N=25). Unpaired t-tests with Welch’s correction were performed between the following groups: control and *Dhc-RNAi* stage 5-stage 9, all p<0.0001 (****); control and *egl-RNAi*, stage 5-stage 9, all p<0.0001 (****); *Dhc-RNAi* and *egl-RNAi*, stage 5, p<0.0001 (****); stage 6, p =0.0369 (*); stage 7, p=0.1825 (n.s); stage 8, p= 0.2122 (n.s); stage 9, p=0.0458 (*). (F-G) The mRNA localizations of *sfGFP-msps 3’UTR* (F) and *CFP-Msps.CDS* (G) via the Cy5-labeled FLAP-X smiFISH probes against sfGFP and CFP, respectively. Both mRNAs show clear oocyte enrichments. All samples with one copy of *maternal αtub-Gal4^[V37]^*. Oocytes are indicated by the yellow arrowheads or yellow brackets. A small (5 µm) (A-C’) or a large (>25 µm) (F-G) Max-intensity Z projections were used to show the mRNA localization. Scale bars, 50 µm. Data are represented as mean ± 95% confidence intervals (D, E).

Next, we demonstrated that the *msps* mRNA accumulation in the oocyte is dependent on the microtubule minus-end-directed motor, dynein. Knockdown of dynein heavy chain (Dhc64C), dynein light intermediate chain (Dlic), or Lis1 all resulted in a “small oocyte” phenotype as previously described (27), and drastically reduces the amount of *msps* mRNA in the oocytes (Figure 3B-3B’ and 3D-3E; Supplementary Figure 3C-3D’). The reduction of *msps* mRNA accumulation becomes more prevalent starting at stage 5, which is consistent with the fact that the *maternal αtub-Gal4[V37]* driver used to express RNAi starts its expression around stage 3∼4 (31). Furthermore, we found that a dynein adaptor Egalitarian (Egl) plays an essential role in localizing *msps* mRNA to the oocyte. Egalitarian is known to be important for linking multiple mRNAs to the dynein motor (34) and is essential for oocyte development and mRNA localization (21, 31). Knockdown of Egl by RNAi driven by *maternal αtub-Gal4^[V37]^* leads to a significantly reduced *msps* mRNA level in the oocyte (Figure 3C-3E), suggesting that Egl is essential for linking the mRNA to the dynein complex. Interestingly, despite the heavily reduced *msps* mRNA in the oocyte, the *egl-RNAi* oocyte is significantly larger than the *Dhc-RNAi* ones, especially in stage 8-9 egg chambers (Supplementary Figure 3E). Noticeably, the ratio of *msps* mRNA average intensity in the oocyte to nurse cells is significantly higher in *egl-RNAi* than in *Dhc-RNAi* (Supplementary Figure 3F), suggesting that the relative abundance of *msps* mRNA could be the key to oocyte determination and growth.

We tested which subregion of *msps* mRNA is required to drive the mRNA accumulation in the oocyte. It turned out that both the coding sequence (CDS) and the 3’ untranslated region (3’-UTR) are sufficient to localize to the mRNA into the oocyte (Figure 3F-3G), while a standard 3’-UTR used in the germline transformation vector results in most mRNAs staying in the nurse cells (Supplementary Figure 3G-3G’). Knockdown of dynein eliminates both the CDS and 3’UTR mRNA accumulation in the oocyte (Supplementary Figure 3H-3I). This is quite different from a single RNA stem-loop structure as the dynein-dependent localization signal in *K10* (transport/localization sequence, TLS), *gurken* (GLS), I-factor Retrotransposon (ILS) and *hairy* (SL1) via the interaction with Egl (34, 60–62). It implies that *msps* mRNA may employ a new localization system for interaction with Egl and its association with the dynein motor.

XMAP215/Msps protein is retained in the oocyte after mRNA accumulation

To examine the localization of Msps protein in the germ line, we generated a CRISPR knock-in line using the newly developed NanoTag epitope, VHH05, (63). We inserted three copies of VHH05 at the C-terminal end of the coding region (Figure 4A; Supplementary Figure 4A-4B). This insertion does not affect the protein functionality as the homozygotes of the CRISPR knock-in line (Msps-3XVHH05) are completely viable and fertile. Coexpression with an EGFP-tagged nanobody specifically reorganizing VHH05 (NbVHH05-EGFP) shows that Msps protein is accumulated in the oocytes by a dynein-dependent mechanism (Figure 4A-4F), similar to *msps* mRNA (Figure 3). This Msps protein enrichment in the oocyte is confirmed using immunostaining with an antibody against Msps (43) (Supplementary Figure 5A), C-terminal tagged Msps-YFP (Supplementary Figure 1E’), and Msps-GFP (64) (Supplementary Figure 5B), and an N-terminal tagged CFP-Msps (65) (Supplementary Figure 5C-5D).

**Figure 4.**
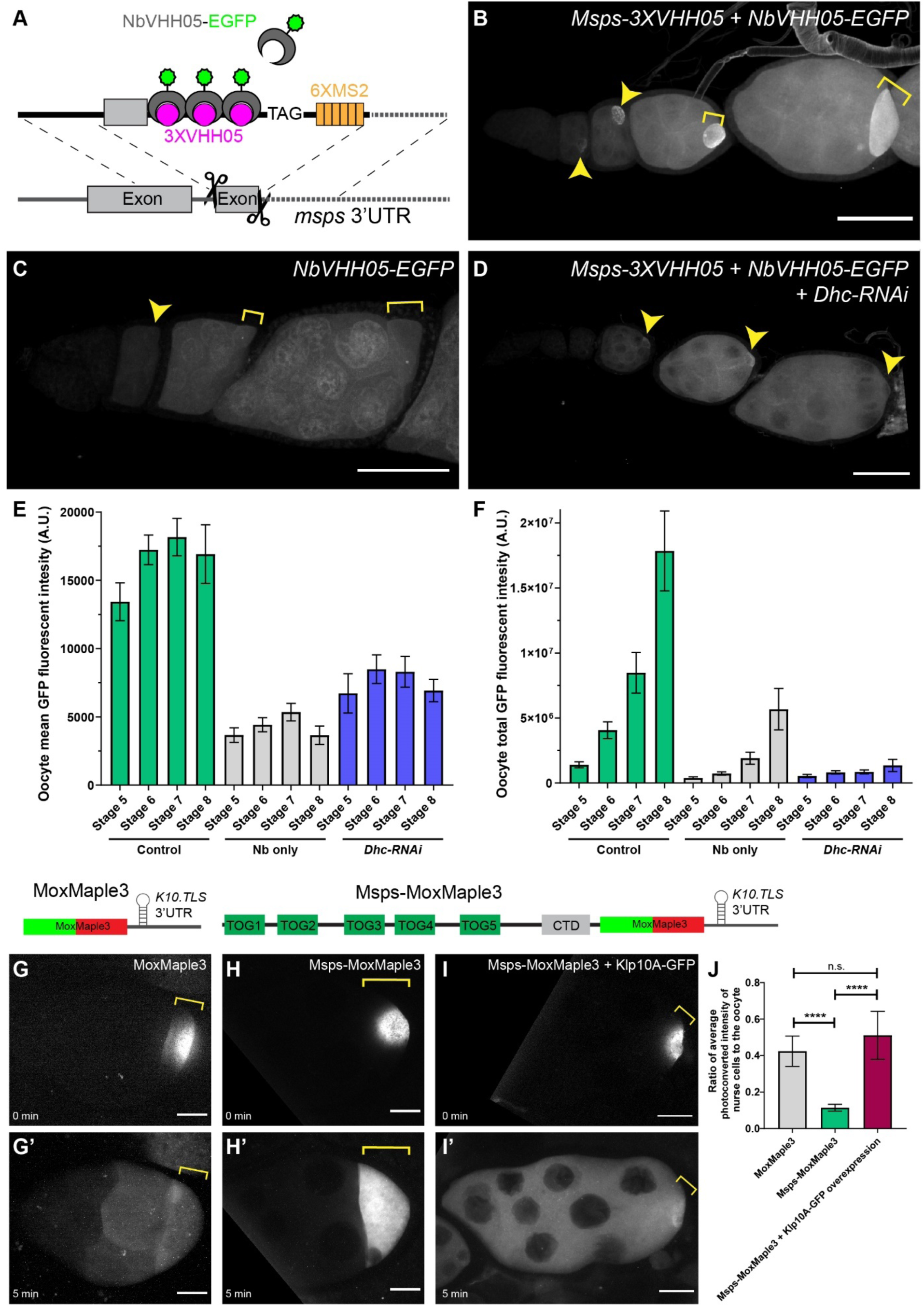
Msps protein is retained in the oocyte. (A) A schematic illustration of labeling Msps protein with the nanotag VHH05. 3 copies of VHH05 are inserted at the C-terminus of Msps CDS via CRISPR-mediated homologous recombination. An EGFP-tagged nanobody specifically recognizing the VHH05 epitope (NbVHH05-EGFP) is expressed to visualize the Msps protein localization. (B-D) Msps protein localization in control (*Msps-3XVHH05 + NbVHH05-EGFP*, B), Nanobody only localization (*NbVHH05-EGFP*, C), and Msps protein localization in *Dhc-RNAi* (*Msps-3XVHH05 + NbVHH05-EGFP + Dhc-RNAi*, D). Msps protein is accumulated in the oocyte in a dynein-dependent manner. (E-F) Average and total fluorescence intensity of NbVHH05-EGFP in control, nanobody only, and *Dhc-RNAi*. (E) The average GFP intensity in control oocytes: stage 5, 13427.4 ± 1380.7 A.U. (N=31); stage 6, 17228.8 ± 1087.8 A.U. (N=37); stage 7, 18158.7 ± 1372.2 A.U. (N=27); stage 8, 16914.8 ± 2143.5 A.U. (N=11). The average GFP intensity in Nanobody (Nb) only: stage 5, 3667.7 ± 530.3 A.U. (N=17); stage 6, 4417.0 ± 514.2 A.U. (N=15); stage 7, 5344.8 ± 643.1A.U. (N=17); stage 8, 3645.7 ± 670.0 A.U. (N=9). The average GFP intensity in *Dhc-RNAi* oocytes: stage 5, 6718.3 ± 1442.3 A.U. (N=18); stage 6, 8487.7 ± 1045.2 A.U. (N=25); stage 7, 8299.3 ± 1129.3 A.U. (N=16); stage 8, 6921.8 ± 821.5 A.U. (N=18). Unpaired t-tests with Welch’s correction were performed between the following groups: control and Nb only, stage 5-stage 8, all p<0.0001 (****); control and *Dhc-RNAi*, stage 5-stage 8, all p<0.0001 (****); Nb only and *Dhc-RNAi*, stage 5-stage 8, all p<0.0001 (****). (F) The total GFP intensity in control oocytes: stage 5, 1401534 ± 231318 A.U. (N=31); stage 6, 4058648 ± 648400 A.U. (N=37); stage 7, 8473610 ± 1557721 A.U. (N=27); stage 8, 17839588 ± 3074993 A.U. (N=11). The total GFP intensity in Nanobody (Nb) only: stage 5, 391365 ± 77689 A.U. (N=17); stage 6, 725633 ± 126531 A.U. (N=15); stage 7, 1906486 ± 461983 A.U. (N=17); stage 8, 5674608 ± 1593674 A.U. (N=9). The total GFP intensity in *Dhc-RNAi* oocytes: stage 5, 533218 ± 123733 A.U. (N=18); stage 6, 816358 ± 122924 A.U. (N=25); stage 7, 856321 ± 153011 A.U. (N=16); stage 8, 1351516 ± 469820. A.U. (N=18). Unpaired t-tests with Welch’s correction were performed between the following groups: control and Nb only, stage 5-stage 8, all p<0.0001 (****); control and *Dhc-RNAi*, stage 5-stage 8, all p<0.0001 (****); Nb only and *Dhc-RNAi*, stage 5, p=0.0496 (*); stage 6, p=0.2864 (n.s.); stage 7, p=0.0002 (***); stage 8, p=0.0002 (***). (G-I’) Photoconverted signals of MoxMaple3 (G-G’), Msps-MoxMaple3 (H-H’), and Msps-MoxMaple3 with Klp10A-GFP overexpression (I-I’) at 0 min (G-I) and 5 min (G’-I’) after local photoconversion in the oocyte. K10 SubregionA containing the dynein-depend localization signal (*K10.TLS*) (60) was inserted at the beginning of the 3’-UTR to ensure the mRNA enrichment in the oocyte. (J) The ratio of the average fluorescence intensity of the red photoconverted signal in nurse cells to the oocyte. For MoxMaple3, the ratio is 0.424 ± 0.08 (N=17); for Msps-MoxMaple3, the ratio is 0.115 ± 0.019 (N=19); for Msps-MoxMaple3 + Klp10A-GFP overexpression, the ratio is 0.511 ± 0.132 (N=10). Unpaired t-tests with Welch’s correction were performed between the following groups: MoxMaple3 and Msps-MoxMaple3, p<0.0001 (****); Msps-MoxMaple3 and Msps-MoxMaple3 + Klp10A-GFP, p<0.0001 (****); MoxMaple3 and Msps-MoxMaple3 + Klp10A-GFP, p= 0.2289 (n.s.). All samples with one copy of *maternal αtub-Gal4^[V37]^*. Oocytes are indicated by the yellow arrowheads or yellow brackets. Scale bars, 50 µm (B-D) and 25 µm (G-I’). Data are represented as mean ± 95% confidence intervals (E, F, J).

**Figure 5.**
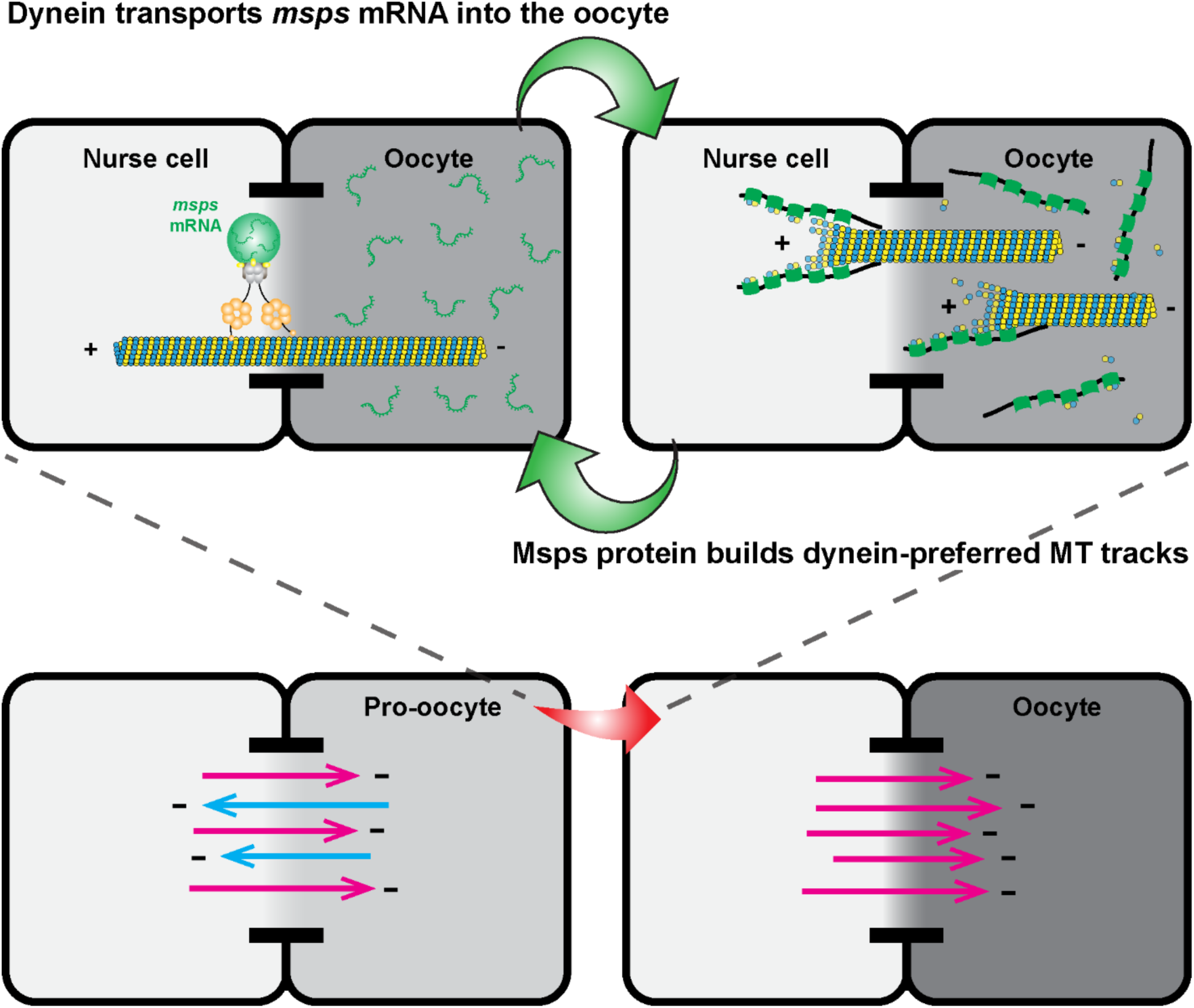
Dynein-Msps positive feedback loop ensures the oocyte determination. The positive feedback loop is driven by the dynein-Msps duo team. Dynein transports *msps* mRNA into the oocyte, while the local translation of Msps protein promotes more microtubule polymerization with plus-ends extending into the nurse cell, which creates dynein-preferred walking tracks. Together, it transforms a slight difference in the number of microtubule minus-ends in the pro-oocyte into a clear cell fate determination of an oocyte. Dynein-dependent minus-end-directed transports into the oocyte and the nurse cell are shown as magenta arrows and blue arrows, respectively.

However, the mRNA accumulation in the oocyte is not sufficient to maintain a high concentration of a protein pool in the oocyte. Adding a dynein-dependent localization signal of *K10* (*K10.TLS*) (60) in the 3’UTR region is sufficient to concentrate the mRNA of MoxMaple3 in the oocyte; however, the MoxMaple3 protein after translation is diffused throughout the whole egg chamber (Supplementary Figure 5E-5E’). Similarly, the translated product of sfGFP-*msps 3’UTR* is not restricted in the oocyte, despite the fact that the mRNA is concentrated in the oocyte (Supplementary Figure 5F-5F’). Furthermore, MoxMaple3-*K10.TLS* photoconverted in the oocyte quickly diffused into nurse cells (Figure 4G-G’ and 4J), indicating that a soluble protein moves easily between the oocyte and interconnected nurse cells via ring canals.

Thus, an active retention mechanism is required to keep a high Msps protein concentration in the oocyte after translation. It is supported by the fact that the photoconverted Msps-MoxMaple-*K10.TLS* remains mostly restricted to the oocyte after photoconversion, unlike MoxMaple-*K10.TLS* (Figure 4G-4H’ and 4J). As Msps interacts with the microtubule lattice and polymerizing plus-ends (42, 43, 45), we speculated that microtubules in the oocyte (Figure 2E and Supplementary Figure 2A) play a role in the retention of Msps protein, preventing its diffusion into nurse cells. To test this hypothesis, we depolymerized microtubules in the oocyte by Klp10A-GFP overexpression (Supplementary Figure 2F-2I). We performed the same photoconversion experiments in these Klp10A-overexpressing ovaries; instead of constraining in the oocyte, Msps-MoxMaple3 diffuses into the interconnected nurse cells from a microtubule-free oocyte (Figure 4I-4J). It indicates that microtubules are essential for retaining Msps protein in the oocyte after mRNA translation.

Altogether, we have shown that: (1) dynein is required for transporting *msps* mRNA into the oocyte; (2) translated Msps protein is retained in the oocyte in a microtubule-dependent manner; (3) high Msps protein concentration in the oocyte is required to assemble microtubules in the oocyte and oocyte-to-nurse cell polarized microtubule network, which favors dynein-dependent transport towards the oocyte. Thus, Msps and dynein form a positive feedback loop to ensure the oocyte has minus-ends-in microtubules than nurse cells (Figure 5; Video 4). Therefore, the Msps-dynein dynamic duo team is essential for oocyte growth and oocyte fate maintenance.

## Discussion

### The positive feedback loop amplifies the initial difference in microtubule polarity

The initiation of the Msps-dynein positive feedback loop requires that one of the cystocytes have a slightly higher concentration of microtubule minus-ends. Among the 16 cystocytes within the *Drosophila* germline cyst, only the two oldest cells, the pro-oocytes, compete to become the oocyte. It has been shown that the pro-oocytes inherited slightly more materials of the fusome, an interconnecting structure enriched with membrane vesicles, actin, and spectrin, than other sister cells (14). Hence, the pro-oocytes are born with more minus-end stabilizing protein, Patronin/CAMSAP, that is recruited to the fusome structure via the spectraplakin protein, Short stop (Shot) (14). Furthermore, the centrosomes are accumulated in the future oocytes after migrating along the fusome, providing further advantages of minus-end nucleating activity over other sister cells (66).

Between the two pro-oocytes, a stochastic difference in microtubule minus-ends may occur after cell division. We propose that the Msps-dynein team can amplify the slight microtubule polarity difference between the two pro-oocytes, as well as between pro-oocyte and other sister nurse cells (Figure 5). Over time, only one winner oocyte is specified and maintained within each 16-cell cyst.

### Egl links *msps* mRNA to the dynein complex and regulates microtubule dynamics

Egl is a dynein adaptor essential for linking mRNAs to the dynein motor. Egl has BicD-binding and Dynein light chain (Dlc)-binding domains at the oppositive ends of the protein and a central mRNA-binding domain (21, 26, 34). Dlc binding facilitates Egl dimerization and thus mRNA binding; in turn, Egl-mRNA binds BicD and gets attached to the dynein motor (32, 35, 67). We found that *msps* mRNA accumulation requires both dynein and the RNA binding Egl, suggesting that Egl links *msps* mRNA to dynein for dynein-dependent transport into the oocyte. Unlike most of the well-known Egl binding mRNA cargoes (e.g., *K10*, *gurken*, *I-factor Retrotransposon*, and *hairy*), *msps* mRNA has more than a single dynein localizing signal, as both of the *msps* CDS mRNA and 3’UTR mRNA are transported into the oocyte in a dynein-dependent manner (Figure 3F-3G; Supplementary Figure 3H-3I). Further studies of narrowing down the dynein-dependent localization signals in the *msps* coding region and the 3’UTR are required to dissect the interaction mechanism. Recently, the positively charged region within the Egl RNA binding domain responsible for binding the localization element of the I factor retrotransposon (ILS) has been mapped (33). Therefore, it would be interesting to test whether this Egl region is also required for interacting with *msps* mRNA.

Previously, it has been shown that the formation of MTOC in the oocyte is disrupted in *BicD* and *egl* mutants (10, 13, 28), indicating that dynein and Egl are actively involved in organizing and maintaining the polarized oocyte microtubule network. Based on our study, the key player in the process is XMAP215/Msps, whose mRNA is transported by dynein to the oocyte via its interaction with Egl and then translated to Msps protein, which in turn stimulates microtubule formation in the oocyte.

### Msps protein retention in the oocyte is microtubule-dependent

Our data suggest that the *msps* mRNA concentration in the oocyte is not sufficient to maintain its protein accumulation in the oocyte, as soluble MoxMaple3 escapes into nurse cells despite that its mRNA is concentrated in the oocyte due to the dynein-dependent localization signal K10.TLS inserted in the 3’-UTR (Figure 4G-G’ and Supplementary Figure 5E-E’). Thus, an active retention mechanism is required to keep produced Msps protein in the oocyte. Based on the fact that Msps binds microtubule lattice and polymerizing plus-ends (42, 43, 45), we hypothesized that the interaction between Msps and microtubules is a key to its retention in the oocyte. We took advantage of the established Klp10A-GFP overexpression line that depolymerizes most of the microtubules in the oocyte and, as a result, decreases the oocyte size (Supplementary Figure 2E-2I). With the Klp10A overexpression, the oocyte-photoconverted Msps protein is seen throughout the entire egg chamber without displaying the tiering pattern (the posterior nurse cells have more photoconverted signal than the anterior nurse cells) we observed in the MoxMaple3 control samples (compare Figure 4I-I’ with 4G-G’). It implies that in the absence of microtubules, Msps protein can freely diffuse between interconnected germline cells. This data supports our hypothesis that Msps is retained within the oocyte via its interaction with microtubules.

However, we cannot rule out the possibility that another protein is needed to mediate Msps and microtubule interaction and, thus, is essential for Msps protein retention. A recent BicD-TurboID protein interactome study suggested that BicD interacts with Msps protein (Graydon Gonsalvez group, unpublished, personal communication). Like other dynein components, BicD protein is accumulated in the oocyte (27, 28), probably after the directed transport along the polarized microtubule network. Therefore, it is also possible that the interaction with BicD enhances Msps protein concentration in the oocyte after local translation.

### The difference in microtubule stability between nurse cells and the oocyte

The dynamics of microtubules in nurse cells and the oocyte are regulated by different mechanisms. The high XMAP215/Msps level promotes a higher level of microtubule polymerization in the oocyte compared to nurse cells (Figure 2; Video 1). At the same time, microtubules in nurse cells are significantly more stable than in the oocyte. Previously, we have shown that the photoconverted microtubules persist in nurse cells for more than 20 min without significant subunit exchange (27), but microtubules photoconverted in the oocytes before stage 10B undergoes very fast depolymerization and repolymerization (68). This explains a very small difference in microtubule amount in nurse cells between control and *msps-RNAi*, as nurse cell microtubules are stable and do not require Msps-dependent polymerization. It raises an interesting possibility that, in addition to the oocyte-concentrated microtubule polymerase XMAP215/Msps, a distinct mechanism stabilizes microtubules in nurse cells, and components of this mechanism are excluded from the oocyte. More studies are needed to characterize the different profiles of the microtubule-associated proteins in nurse cells and the oocyte, and further understand the differential regulations of these microtubule populations.

Altogether, we have identified a positive feedback loop involving the microtubule polymerase, XMAP215/Msps, and the microtubule minus-end-directed motor, dynein. We propose that the Msps-dynein team amplifies a small initial difference in microtubule density and polarity between sister cells and transforms it into a uniformly polarized microtubule network to ensure oocyte differentiation and maintenance.

## Materials and methods

### *Drosophila* husbandry and maintenance

Fly stocks and crosses were kept on standard cornmeal food (Nutri-Fly Bloomington Formulation, Genesee, Cat #: 66–121) supplemented with active dry yeast in the 24∼25C incubator. Following flies were used in this study: *mat αtub-Gal4^[V37]^* (III, Bloomington *Drosophila* Stock Center #7063); *nos-Gal4^[VP16]^* (III) (51, 69); *UAS-msps-RNAi* (HMS01906, attP40, II, Bloomington *Drosophila* Stock Center # 38990, targeting Msps CDS 5001-5021 nt, 5’-CTGCGCGACTATGAAGAAATA-3’); *ubi-EB1-GFP* (III) (from Dr. Steve Roger, the University of North Carolina at Chapel Hill) (50, 54–56); *UASp-EB1-GFP* (II, from Dr. Antoine Guichet) (50, 70); *ubi-GFP-Pav* (II) (50, 71); *mat αtub67C-EMTB-3XGFP-sqh 3’UTR* (attP40, II, an unpublished gift from Dr. Yu-Chiun Wang, RIKEN Center for Biosystems Dynamics Research) (27); *UASp-Klp10A-GFP-SspB* (II) (59); *UAS-Dhc64C-RNAi* (TRiP.GL00543, attP40, II, Bloomington *Drosophila* Stock Center #36583, targeting DHC64C CDS 10044–10,064 nt, 5’-TCGAGAGAAGATGAAGTCCAA-3’) (27, 54, 55); *UAS-Dlic-RNAi* (targeting Dlic 3’UTR 401–421 nt, 5’-AGAAATTTAACAAAAAAAAAA –3’, III, inserted at attP-9A 75A10 site) (27); *UAS-Lis1-RNAi* (II, from Dr. Graydon Gonsalvez, Augusta University, targeting Lis1 CDS 1197– 1217 nt, 5’-TAGCGTAGATCAAACAGTAAA-3’) (27, 72); *UAS-Egl-RNAi* (TRiP.GL01170, attP2, III, Bloomington *Drosophila* Stock Center #43550, targeting Egl CDS 1590-1610 nt, 5’-CACGGTGATAGCGAATGTCAA-3’); *UASp-CFP-Msps.CDS* (from Dr. Timothy Megraw, Florida State University) (Zheng et al., 2020); *Tub-PBac* (Bloomington *Drosophila* Stock Center #8285); *UASt-NbVHH05-EGFP* (attP40, II, Bloomington *Drosophila* stock center # 94008) (63); *UAS-Rhi-RNAi* (From Dr. Zhao Zhang, Duke University School of Medicine) (73); *ubi-Msps.CDS-GFP* (From Dr. Jordan Raff, University of Oxford) (Lee et al., 2001). The following transgenic fly stocks were generated in this study using PhiC31-mediated integration (BestGene Inc.): *UASp-sfGFP-msps 3’UTR and UASp-sfGFP-K10CT 3’UTR* (inserted at 89E11, III); *UASp-MoxMaple3-K10subregionA 3’UTR* and *UASp-Msps (RNA-resistant)-MoxMaple3-K10subregionA 3’UTR* (inserted at 75A10, III); *UASp-Msps (RNA-resistant)-YFP-BLID-K10subregionA 3’UTR* (inserted at 55C4, II).

### CRISPR Knock-In to create *msps-3XVHH05-6XMS2*

Two small gRNA (#1-GGGGTATTTCAATCAGAAGC; #2-ACGGGAAGCGCACAGTTTAT) targeting *msps* genomic region were synthesized and inserted into the pCFD5 vector (74) (Addgene, Plasmid #73914) via BbsI digestion. 6XMS2 were amplified from the pSL-MS2-6X (Addgene, plasmid #27118) (75) and inserted into the pScarlessHD-C-3xVHH05-DsRed (Addgene, Plasmid #171580) (63) via InFusion cloning (Takara Bio) to create the vector of pScarlessHD-C-3xVHH05-6XMS2-DsRed. ∼1 kb 5’ homology arm and ∼1 kb 3’ homology arm of *msps* genomic region were amplified from the genomic DNA (Sigma, Extract-N-Amp™ Tissue PCR Kit), mutated to be insensitive to gRNAs, and inserted into the pScarlessHD-C-3xVHH05-6XMS2-DsRed vector via InFusion cloning (Takara Bio). The DNA plasmid of pScarlessHD-5’ *msps* homology arm-C-3xVHH05-6XMS2-DsRed-3’ *msps* homology arm was co-injected with pCFD5-gRNA#1 and pCFD5-gRNA#2 by BestGene. Flies with fluorescent red eyes were selected and crossed with *Tub-PBac* flies to remove the DsRed region by PBac transposase. The final *msps-3XVHH05-6XMS2* line was verified using genomic PCR amplification and sequencing.

### Plasmid constructs

#### -pUASp-attB-sfGFP-msps 3’UTR

*msps* 3’UTR were amplified from the genomic DNA and inserted into the pUASp-attB-ΔK10 (the original C-terminus (CT) of K10 3’UTR and terminator region was replaced with *Drosophila* α-tubulin terminator and polyA signal, a kind gift from Dr. Paul Schedl, Princeton University) (76) via EcoRI(5’) and KpnI(3’) to create pUASp-attB-msps 3’UTR. sfGFP was amplified by PCR and inserted into the pUASp-attB-msps’3UTR via XbaI (5’) and EcoRI (3’) to create pUASp-attB-sfGFP-msps 3’UTR.

#### -pUASp-attB-sfGFP-K10CT 3’UTR

The K10CT 3’UTR (973-1489 nt, after HpaI site, without the subregion A or TLS that is required for oocyte transport and localization) (60) were amplified by PCR from the pUASp vector (77) and inserted into the pUASp-attB-sfGFP-msps 3’UTR via EcoRI(5’) and KpnI(3’) to replace the *msps* 3’UTR.

#### -pUASp-attB-MoxMaple3-K10SubregionA 3’UTR

The MoxMaple3 were amplified from pUASp-Mito-MoxMaple3 construct (50) by PCR and inserted into pUASp-attB via SpeI(5’) and XbaI(3’). A small fragment of K10 3’UTR (Subregion A containing TLS, AGGCCTTAGATTACACCACTTGATTGTATTTTTAAATTAATTCTTAAAAACTACAAATTAAGATCACTCTGTGAACGTGTGCTCGATGGTG)(60) was synthesized and inserted into the pUASp-attB-MoxMaple3 via PspXI single digestion to ensure the oocyte enrichment of the *MoxMaple3* mRNA.

#### -pUASp-attB -Msps (RNA-resistant)-MoxMaple3-K10subregionA 3’UTR

Two DNA fragments carrying overlapping silent mutations of Msps protein isoform C (7225-7245 nt, TTAAGAGATTACGAGGAGATT, corresponding to 1636-1642 residues of LRDYEEI, RNAi-resistant to *UAS-msps-RNAi* TRiP line HMS01906 that targets 5’-CTGCGCGACTATGAAGAAATA-3’) were amplified from the pIZ-Msps-GFP construct (a kind gift from Dr. Steve Roger, the University of North Carolina at Chapel Hill) (43) and inserted back into pIZ-Msps-GFP digested with SspI(5’) and XhoI(3’) using Infusion ligation (Takara Bio) to create pIZ-Msps (RNAi-resistant)-GFP. Msps (RNAi-resistant) was then subcloned into pUASp-attB-MoxMaple3 construct via NotI(5’) and SpeI(3’) to create pUASp-attB-Msps (RNAi-resistant)-MoxMaple3. A small fragment of K10 3’UTR (Subregion A containing TLS, GGCCTTAGATTACACCACTTGATTGTATTTTTAAATTAATTCTTAAAAACTACAAATTAAGATCACTCTGTGAACGTGTGCTCGATGGTG) (60) was synthesized and inserted into the pUASp-attB-Msps (RNAi-resistant)-MoxMaple3 via PspXI single digestion to ensure the oocyte enrichment of the *msps (RNAi-resistant)-MoxMaple3* mRNA.

#### -pUASp-Msps (RNA-resistant)-YFP-BLID-K10subregionA 3’UTR

YFP-BLID (sensitive to blue light; delete the last three amino acids from AsLOV2, with I532A mutation and RRRG degron) was amplified by PCR from pBMN-HA-YFP-LOV24 (Addgene, Plasmid #49570) (78) and inserted into the pUASp-attB vector via BamHI(5’) and XbaI(3’). Msps (RNAi-resistant) fragment was subcloned from pUASp-attB-Msps (RNAi-resistant)-MoxMaple3 into pUASp-attB-YFP-BLID via NotI (5’) and SpeI (3’). A small fragment of K10 3’UTR (Subregion A containing TLS, AGGCCTTAGATTACACCACTTGATTGTATTTTTAAATTAATTCTTAAAAACTACAAATTAAGATCACTCTGTGAACGTGTGCTCATGGTG) (60) was synthesized and inserted into the pUASp-attB-Msps (RNA-resistant)-YFP-BLID via PspXI single digestion to ensure the oocyte enrichment of *msps (RNAi-resistant)-YFP-BLID* mRNA.

### Live imaging of *Drosophila* egg chambers

Young female adults were mated with several male flies and fed with active dry yeast for 16∼18 hours before dissection. The ovaries were dissected in Halocarbon oil 700 (Sigma-Aldrich, Cat# H8898) as previously described (27, 50, 59, 68, 79). Freshly dissected samples were imaged on a Nikon W1 spinning disk confocal microscope (Yokogawa CSU with pinhole size 50 µm) with a Hamamatsu ORCA-Fusion Digital CMOS Camera, and a 40X 1.25 N.A. silicone oil lens, controlled by Nikon Elements software.

### Anti-Orb Immunostaining in *Drosophila* egg chambers

A standard fixation and immunostaining protocol was described previously (27, 50, 59, 68, 69, 79). Young female adults were mated with several male flies and fed with active dry yeast for 16∼18 hours before dissection. Ovaries were dissected in 1X PBS and fixed with 4% EM-grade formaldehyde (Electron Microscopy Sciences 16% Paraformaldehyde Aqueous Solution, Fisher Scientific, Cat# 50-980-487) in 1X PBS +0.1% Triton X-100 for 20 min on a rotator; washed with 1X PBTB (1XPBS +0.1% Triton X-100+0.2% BSA) five times for 10 min each time, and blocked in 5% (vol/vol) normal goat serum-containing 1X PBTB for 1h at RT; stained with the primary mouse monoclonal anti-Orb antibody (Orb 4H8, Developmental Studies Hybridoma Bank, supernatant, 1:5) at 4 °C overnight and with the secondary FITC-conjugated or TRITC-conjugated anti-mouse secondary antibody (Jackson ImmunoResearch Laboratories, Inc; Cat# 115-095-062 and Cat# 115-025-003) at 10 µg/ml at room temperature (24∼25 °C) for 4 hours; stained with rhodamine-conjugated or Alexa Fluor633-conjugated phalloidin (0.2 µg/ml), and DAPI (1 µg/mL) for >1 hour at room temperature before mounting, and imaged on a Nikon A1plus scanning confocal microscope with a GaAsP detector and a 20X 0.75 N.A. lens using Galvano scanning, controlled by Nikon Elements software. Z-stack images were acquired every 1 µm/step.

### Microtubule staining in *Drosophila* egg chambers

Ovaries were dissected in 1X Brinkley Renaturing Buffer 80 (BRB80, 80 mM piperazine-N,N’-bis(2-ethanesulfonic acid) [PIPES], 1 mM MgCl2, 1 mM EGTA, pH 6.8) and fixed in 8% EM-grade formaldehyde (Electron Microscopy Sciences 16% Paraformaldehyde Aqueous Solution, Fisher Scientific, Cat# 50-980-487) +1X BRB80 +0.1% Triton X-100 for 20 min on a rotator; briefly washed with 1X PBTB (1X PBS + 0.1% Triton X-100 +0.2% BSA) five times for 10 min each time, and blocked in 5% (vol/vol) normal goat serum-containing 1X PBTB for 1 hour at room temperature; stained with CoraLitePlus 488conjugated β-tubulin monoclonal antibody (ProteinTech, Cat# CL488-66240, CloneNo. 1D4A4) or CoraLite594-conjugated β-tubulin monoclonal antibody (Proteintech, Cat# CL594-66240 CloneNo. 1D4A4) 1:100 at 4 °C overnight; samples were stained rhodamine-conjugated or Alexa Fluor633-conjugated phalloidin (0.2 µg/ml), and DAPI (1 µg/mL) for 1 hour before mounting. Samples were imaged using a Nikon W1 spinning disk confocal microscope (Yokogawa CSU with pinhole size 50 µm) with a Hamamatsu ORCA-Fusion Digital CMOS Camera and a 100X 1.35 N.A. silicone oil lens, controlled by Nikon Elements software. Images were acquired every 0.5 µm/step in z stacks.

Ovaries from flies expressing maternal αtub67C-EMTB-3XGFP-sqh 3’UTR were dissected in 1X BRB80 buffer and fixed in 8% EM-grade formaldehyde +1X BRB80 +0.1% Triton X-100 for 20 min on a rotator; briefly washed with 1X PBTB five times for 10 min each time, and stained rhodamine-conjugated phalloidin (0.2 µg/ml) and DAPI (1 µg/mL) for 1 hour before mounting. Samples were imaged using a Nikon W1 spinning disk confocal microscope (Yokogawa CSU with pinhole size 50 µm) with a Hamamatsu ORCA-Fusion Digital CMOS Camera and a 100X 1.35 N.A. silicone oil lens, controlled by Nikon Elements software. Images were acquired every 0.5 µm/step in z stacks.

### Single-molecule inexpensive fluorescence *in-situ* hybridization (smiFISH)

smiFISH protocol was based on (80–82) with some small modifications. 20 base-long DNA probes complementary to the mRNA of *msps* 3’UTR, sfGFP, CFP, or MoxMaple3 with 3’ FLAP-X complementary probe (5’-CCTCCTAAGTTTCGAGCTGGACTCAGTG-3’) were designed using LGC Biosearch Technologies’ Stellaris RNA FISH Probe Designer (masking level 5, minimal spacing 2 bases). 25 probes specific to *msps* 3’UTR mRNA and 10 probes specific to sfGFP, CFP, and MoxMaple3 mRNAs were ordered from ThermoFisher (25 nmol synthesis scale, standard desalting) (the probe sequences are listed in Supplemental Table 1) and diluted to 100 μM in nuclease-free H2O. Probes were mixed at equal molar ratios (to a mixed probe concentration of 100 μM) and stored at -20°C. Fluorescently labeled Flap-X probe with 5’ and 3’ Cy5 modifications (/5Cy5/CACTGAGTCCAGCTCGAAACTTAGGAGG-/3Cy5Sp/) was ordered from IDT (100 nmol synthesis scale, HPLC purified), diluted in nuclease-free H2O to a concentration of 100 μM, and stored at −20 °C in aliquots. mRNA-FLAP-X binding Probes and fluorescent Flap-X probes were annealed by mixing 2 μl of mRNA-FLAP-X mixed probe (100 μM mixed concentration), 2.5 μl of Cy5-FlapX-binding probe (100 μM), 5 μl of New England Biolabs Buffer 3 (1X composition: 100mM NaCl, 50mM Tris-HCl, 10mM MgCl2, 1mM DTT, pH 7.9), and 40.5 μl nuclease-free H2O, incubated at 85°C for 3 min, 65°C for 3 min, and 25°C for 5 min in a PCR machine, and stored at −20 °C.

Ovaries were (1) dissected in 1X PBS and fixed in 4% EM-grade formaldehyde (Electron Microscopy Sciences 16% Paraformaldehyde Aqueous Solution, Fisher Scientific, Cat#50-980-487) in 1X PBST (1XPBS +0.1% Triton X-100 in nuclease-free H_2_O) for 20 min on a rotator at room temperature; (2) washed three times in 1X PBST for 5 min each at room temperature; (3) exchanged into a 1:1 volume mixture of 1X PBST and smiFISH Wash Buffer [10% 20X SSC (20X SSC: 0.3M sodium citrate, 3M NaCl, in nuclease-free H_2_O, pH7.0), 10% deionized formamide, in nuclease-free water], and incubated at room temperature for 10 min; (4) washed two times in smiFISH Wash Buffer for 10 min each at room temperature; (5) incubated at 37°C in smiFISH Wash Buffer for 30 min; (6) incubated with 2 µl annealed probe diluted in 37°C prewarmed 500 µl smiFISH Hybridization Buffer (10% dextran sulfate, 10% 20X SSC, 10% deinionzed formamide, in nuclease-free water) overnight (>16 hours) at 37°C in dark; (7) diluted with 37°C 500 μl of smiFISH Wash Buffer; (8) washed three times in smiFISH Wash Buffer for 10 min each at 37°C; (9) incubated at room temperature in a 1:1 volume mixture of 1X PBST and smiFISH Wash Buffer for 10 min; (10) incubated in 1X PBST with rhodamine-conjugated phalloidin (0.2 µg/ml) and DAPI (1 µg/mL) at the overnight smiFISH Hybridization Buffer for 1 hour; (11) washed two times in 1X PBST at room temperature for 10 min each before mounting.

### Measurement of nurse cell and oocyte size

Z stacks of triple color images of the ovarioles stained with DAPI, anti-Orb antibody, and rhodamine conjugated-phalloidin staining were acquired on a Nikon A1plus scanning confocal microscope with a GaAsP detector and a 20X 0.75 N.A. lens using Galvano scanning. Egg chamber stages characterization was as previously described (50, 83). Nurse cell area and oocyte area were specified (at the largest cross-section) and measured by manual polygon selection (area size) in FIJI.

### Quantification of smiFISH staining

Z stacks of triple color images of the ovarioles stained with DAPI, rhodamine conjugated-phalloidin, and Cy5-labeled FLAP-X smiFISH probe staining were acquired on a Nikon W1 spinning disk confocal microscope (Yokogawa CSU with pinhole size 50 µm) with a Hamamatsu ORCA-Fusion Digital CMOS Camera, and a 40X 1.25 N.A. silicone oil lens, at every 0.5 µm/step. Egg chamber stages characterization was as previously described (50, 83). A sum Z-projection of a total 2.5 µm z-stack image (6 z slices in total) of each egg chamber was created by FIJI (Image>Stacks>Z projection>Sum slices), and cell area and fluorescence intensity were measured by manual polygon selection in FIJI.

### Quantification of microtubule staining in *Drosophila* egg chambers

Z stacks of triple color images of the ovarioles stained with DAPI, rhodamine conjugated-phalloidin, FITC-conjugated β-tubulin staining or EMTB-3XGFP labeling were acquired on a Nikon W1 spinning disk confocal microscope (Yokogawa CSU with pinhole size 50 µm) with a Hamamatsu ORCA-Fusion Digital CMOS Camera, with a 100X 1.35 N.A. silicone oil lens, at every 0.5 µm/step. Egg chamber stages characterization was as previously described (50, 83). A sum Z-projection of a total 2.5 µm z-stack image (6 z slices in total) of a stage 6-7 egg chamber was created by FIJI (Image>Stacks>Z projection>Sum slices), and cell area and fluorescence intensity were measured by manual polygon selection in FIJI.

### EB1 comet tracking in *Drosophila* egg chambers

*ubi-EB1-GFP* time-lapse movies of stage 6-7 egg chambers in either control or *msps-RNAi* were acquired on a Nikon W1 spinning disk confocal microscope (Yokogawa CSU with pinhole size 50 µm) with a Hamamatsu ORCA-Fusion Digital CMOS Camera, and a 40X 1.25 N.A. silicone oil lens, at a frame rate of every 2 seconds for a total of 2 min, controlled by Nikon Elements software. Images were processed in Fiji and analyzed DiaTrack 3.04 Pro (84), with a maximum particle jump distance of 0.57 μm/s, a minimum speed limit of 0.01 μm/s, and a minimum lifetime of 6 sec. The sum of the length of EB1-GFP tracks was considered to be the total length of newly polymerized microtubules. For each sample, both the number of EB1 comets and the length of newly polymerized microtubules are normalized by the cell area size to get the EB1 density (#/µm^2^) and the new microtubule length in 2-min movie (µm/µm^2^).

### Quantification of NbVHH05 staining

Samples of (1) yw/w; UASt-NbVHH05-EGFP/+; Msps-3XVHH05/mat αtub-Gal4^[V37]^, UAS-Rhi-RNAi (control); (2) yw/w; UASt-NbVHH05-EGFP/+; mat αtub-Gal4^[V37]^, UAS-Rhi-RNAi/+ (Nb only); (3) yw/w; UASt-NbVHH05-EGFP/UAS-Dhc-RNAi; Msps-3XVHH05/mat αtub-Gal4^[V37]^, UAS-Rhi-RNAi (Dhc-RNAi) were dissected in 1X PBS and fixed with 4% EM-grade formaldehyde in 1X PBS +0.1% Triton X-100 for 20 min on a rotator; washed with 1X PBTB (1X PBS + 0.1% Triton X-100 +0.2% BSA) five times for 10 min each time, and stained with rhodamine-conjugated phalloidin (0.2 µg/ml) and DAPI (1 µg/mL) for >1 hour before mounting. Samples were imaged on a Nikon W1 spinning disk confocal microscope (Yokogawa CSU with pinhole size 50 µm) with a Hamamatsu ORCA-Fusion Digital CMOS Camera, with a 40X 1.25 N.A. silicone oil lens, at every 0.5 µm/step. Egg chamber stages characterization was as previously described (50, 83). A sum Z-projection of a total 5 µm z-stack image (11 z slices in total) of each egg chamber was created by FIJI (Image>Stacks>Z projection>Sum slices), and cell area and fluorescence intensity were measured by manual polygon selection in FIJI.

### Photoconversion of MoxMaple3 and Msps-MoxMaple3

Young mated female adults of (1) *yw; mat αtub-Gal4^[V37]^*/*UASp-MoxMaple3-K10.TLS* and (2) yw; *mat αtub-Gal4^[V37]^/UASp-Msps-MoxMaple3-K10.TLS* were dissected in Halocarbon oil 700, as described above. Freshly dissected samples were imaged on a Nikon W1 spinning disk confocal microscope (Yokogawa CSU with pinhole size 50 µm) with a Hamamatsu ORCA-Fusion Digital CMOS Camera and a 40X 1.25 N.A. silicone oil lens. MoxMaple3 was photoconverted within a circle of ∼12 µm diameter by 405 nm light controlled by Mightex Polygon DMD Illuminators. The photoconverted signal was acquired at 0.5 µm/step in Z-stack images 5 min after photoconversion. Sum slices of the Z projection were used to calculate the ratio of the average intensity of the photoconverted signal in nurse cells to the oocyte.

### Anti-Msps antibody staining

The rabbit crude serum containing the rabbit polyclonal antibody against *Drosophila* Msps TOG2 domain (a kind gift from Dr. Steve Rogers, UNC at Chapel Hill) (43) was pre-adsorbed twice with wild-type ovaries (fixed with 4% EM-grade formaldehyde, but not blocked) at 1:10 dilution at 4°C for overnight to reduce the non-specific background; the pre-absorbed anti-Msps antibody was then diluted 1:5 in overnight staining (final dilution at 1:50) at 4°C; FITC-conjugated anti-rabbit secondary antibody (Jackson ImmunoResearch Laboratories, Inc; Cat#111-095-003) was used at 10 µg/ml at room temperature for 4 hours before washing and mounting. Samples were imaged on a Nikon W1 spinning disk confocal microscope (Yokogawa CSU with pinhole size 50 µm) with a Hamamatsu ORCA-Fusion Digital CMOS Camera and a 40X 1.25 N.A. silicone oil lens at every 0.5 µm/step.

### Statistical analysis

The figures plots show either percentage of phenotypes or mean values, as indicated in figure legends. Error bars represent 95% confidence intervals and N stands for sample numbers examined in each assay. Statistical analysis was performed in GraphPad Prism 8.0.2, and p values with levels of significance are listed in figure legends.

## Data and materials availability

All the data are included in the article and supporting information. Request for materials (DNA plasmids, fly lines, etc.) generated in this study can be made to the corresponding author Vladimir I. Gelfand.

## Author contributions

W.L. and V.I.G. planned and designed the research. W.L., M.L., and V.I.G. conducted experiments and data analysis; W.L. and V.I.G. wrote the manuscript.

## Acknowledgments

We thank many colleagues who generously contributed reagents for this work: Dr. Steve Rogers (UNC at Chapel Hill), Dr. Antoine Guichet (CNRS, Institut Jacques Monod), Dr. David Glover (Caltech), Dr. Yu-Chiun Wang (RIKEN Center for Biosystems Dynamics Research, Japan), Dr. Graydon Gonsalvez (Augusta University), Dr. Timothy Megraw (Florida State University), Dr. Jordan Raff (University of Oxford), Dr. Zhao Zhang (Duke University School of Medicine), Dr. Paul Schedl (Princeton University); and Bloomington *Drosophila* Stock Center (supported by National Institutes of Health grant P40OD018537) for fly stocks. The Orb 4H8 monoclonal antibody developed by Dr. Paul D. Schedl’s group at Princeton University was obtained from the Developmental Studies Hybridoma Bank, created by the NICHD of the NIH and maintained at The University of Iowa. We thank Northwestern Center for Advanced Microscopy & Nikon Imaging Center for imaging assistance and Northwestern Sanger Sequencing Facility for sequencing services. We thank all the Gelfand laboratory members for their support, discussion, and suggestions. W. Lu dedicates this work to the memory of her grandfather, Yuedong Lu, who deceased in December 2022. Research reported in this study was supported by the National Institute of General Medical Sciences grant R35GM131752 to V.I.G.

## Video legends

**Video 1. EB1-GFP comets in control, related to Figure 2.** Polymerizing microtubule plus-ends are labeled with *ubi-EB1-GFP*, and ring canals are labeled with *ubi-GFP-Pav/kinesin-6*. Compared to the neighboring nurse cells, the oocyte has clear enrichment of EB1-GFP comets. Scale bar, 25 µm.

**Video 2. EB1-GFP comets in a control nurse cell-to-oocyte ring canal, related to Figure 2**. Polymerizing microtubule plus-ends are labeled with *UASp-EB1-GFP driven by nos-Gal4*^[VP16]^, and ring canals are labeled with *ubi-GFP-Pav/kinesin-6*. More microtubule plus-ends are growing from the oocyte into the nurse cell. Scale bar, 5 µm.

**Video 3. EB1-GFP comets in *msps-RNAi*, related to Figure 2**. Polymerizing microtubule plus-ends are labeled with ubi-EB1-GFP, and ring canals are labeled with ubi-GFP-Pav/kinesin-6 in the germline *msps-RNAi* driven by *mat αtub-Gal4^[V37]^*. No visible EB1-GFP comets can be seen in the oocyte or nurse cells. To note, the EB1-GFP comets are still visible in the follicle cells, as the germline knockdown of *msps* does not affect somatic follicle cells. Scale bar, 25 µm.

**Video 4. A positive feedback mechanism composed of microtubule minus-end directed motor dynein and microtubule polymerase XMAP215/Msps, related to Figure 5.** Dynein transports more msps mRNA into the oocyte along the polarized oocyte-to-nurse cell microtubule network (the top panel), while translated Msps protein builds more dynein-preferred microtubule tracks from the oocyte into the nurse cell (the bottom panel).

---

**Supplementary Figure 1.**
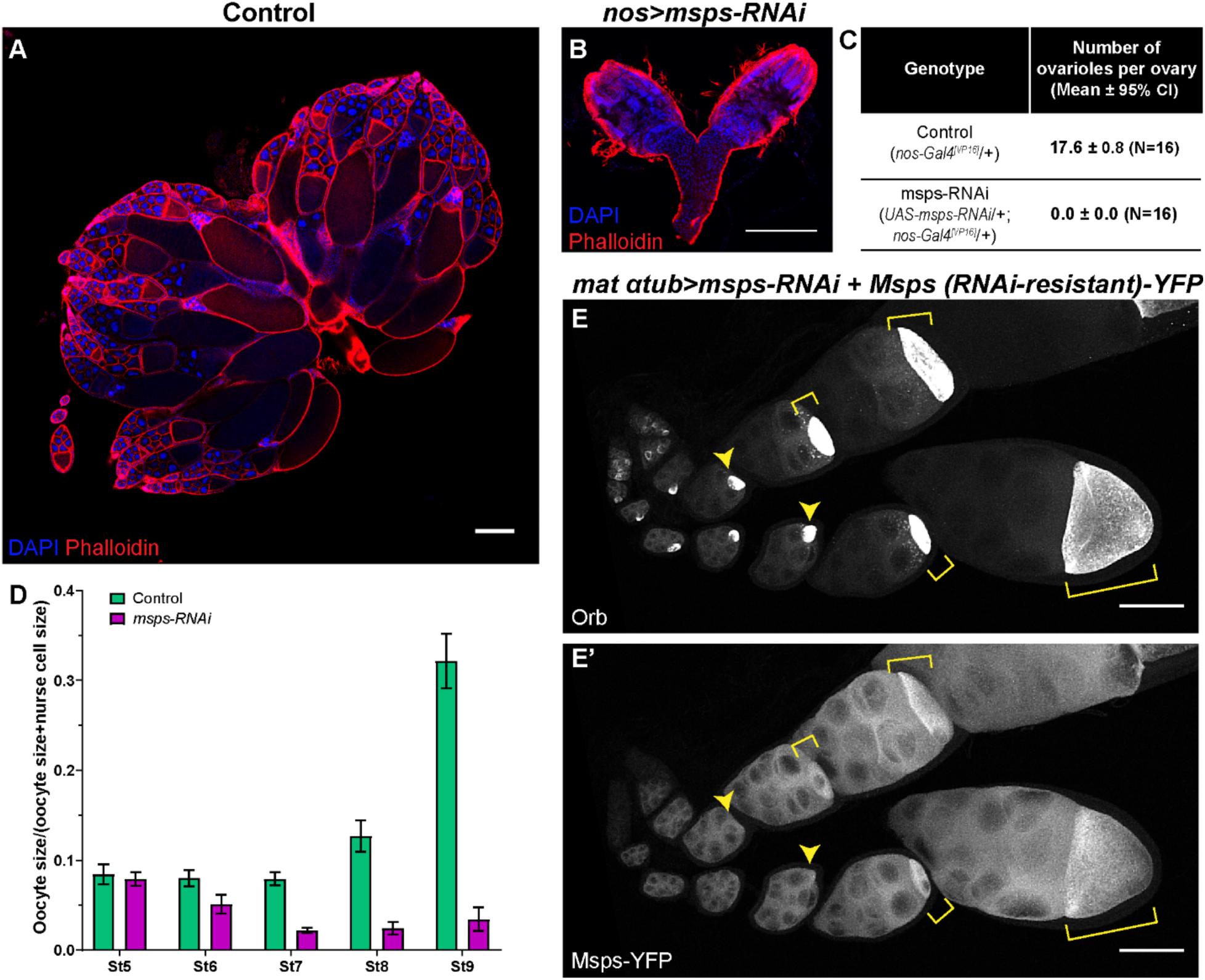
Msps is required for oocyte specification and maintenance. (A-B) Representative ovary images of control (A) and *msps-RNAi* driven by the early germline-specific *nos-Gal4^[VP16]^* (B). Scale bars, 200 µm. (C) The summary of ovariole numbers per ovary in control and *nos>msps-RNAi* (mean ± 95% confidence intervals). Early germline knockdown of *msps* caused a complete germless phenotype. (D) The fraction of oocyte size in germline cells, calculated as oocyte size / (oocyte size+ nurse cell size). The fraction of oocyte size in control: stage 5, 0.084 ± 0.011 (N=66); stage 6, 0.080 ± 0.009 (N=59); stage 7, 0.079 ± 0.007 (N=44); stage 8, 0.127 ± 0.017 (N=25); stage 9, 0.322 ± 0.030 (N=20). The fraction of oocyte size in *msps-RNAi*: stage 5, 0.079 ± 0.008 (N=89); stage 6, 0.051 ± 0.010 (N=79); stage 7, 0.022 ± 0.003 (N=68); stage 8, 0.024 ± 0.007 (N=35); stage 9, 0.034 ± 0.013 (N=27). Data are presented as mean ± 95% confidence intervals. Unpaired t-tests with Welch’s correction were performed between control and *msps-RNAi* oocytes: stage 5, p= 0.4149 (n.s.); stage 6, p <0.0001 (****); stage 7, p <0.0001 (****); stage 8, p <0.0001 (****); stage 9, p <0.0001 (****). (E-E’) Representative images of Orb staining and Msps-YFP labeling in rescued *msps-RNAi* samples. Oocytes are indicated by the yellow arrowheads or yellow brackets. Scale bars, 50 µm.

**Supplementary Figure 2.**
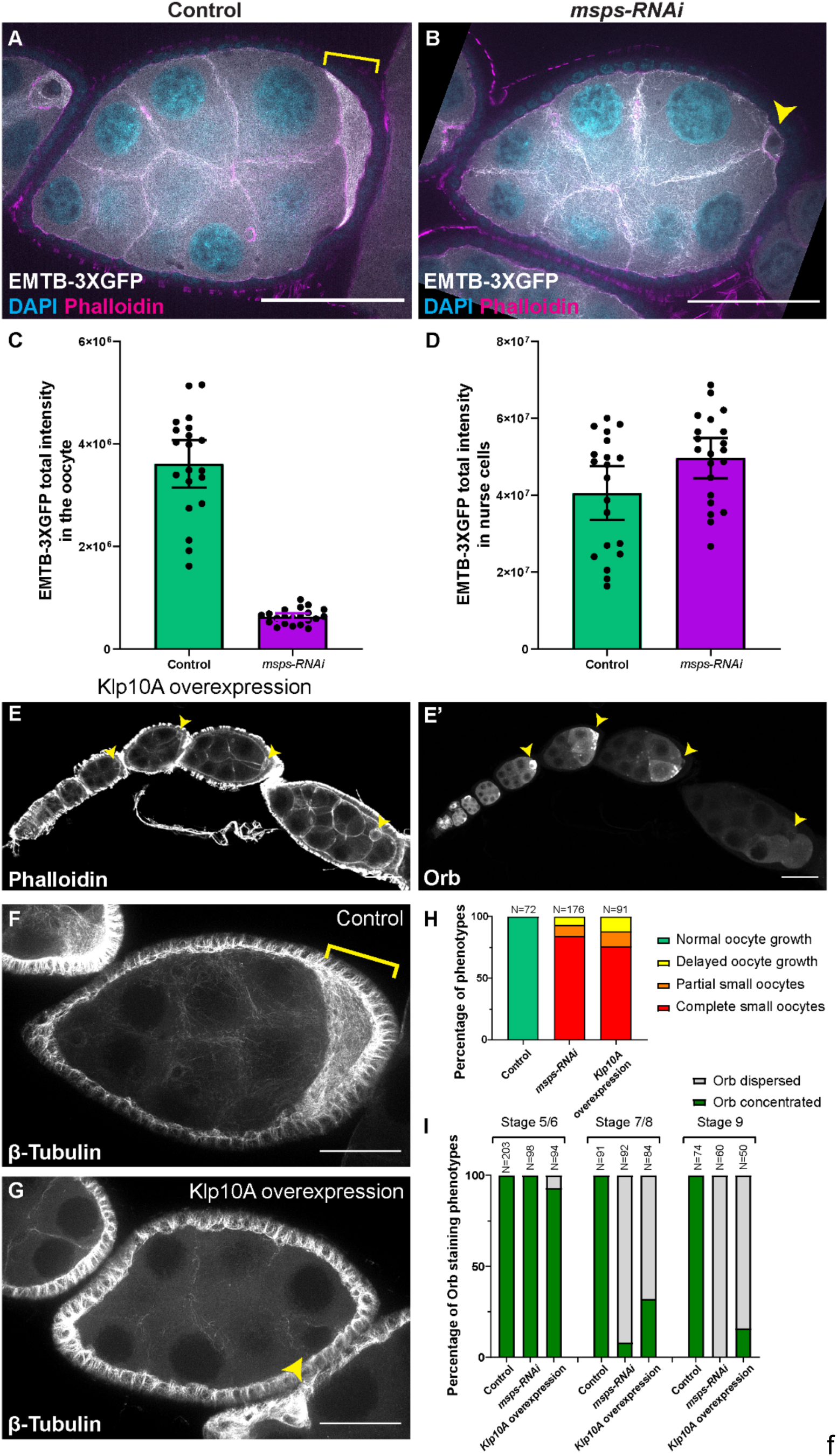
Msps is required for microtubule formation in the oocyte. (A-B) Microtubules labeled with the microtubule-binding domain of human MAP7/Enscosin (EMTB), tagged with 3 copies of GFP in control (A) and *msps-RNAi* (B). Both samples carried one copy of *maternal αtub-Gal4^[V37]^*. Sum-slice projections of a total of 2.5 µm Z-stack images (0.5 µm per step) were used to show the overall microtubule distribution. Oocytes are highlighted with either the yellow bracket (A) or the yellow arrowhead (B). Scale bars, 50 µm. (C-D) Quantification of total EMBT-3XGFP fluorescence intensity in oocytes (C) and nurse cells of stage 6-7 egg chambers (D). Data are represented as mean ± 95% confidence intervals. EMTB-3XGFP total intensity in control stage 6-7 oocytes, 3616040 ± 463397 A.U. (N=20); in *msps-RNAi* stage 6-7 oocytes, 630469 ± 67656 A.U. (N=21). An unpaired t-test with Welch’s correction was performed between control and *msps-RNAi* oocytes, p<0.0001 (****). EMTB-3XGFP total intensity in control stage 6-7 nurse cells, 40565248 ± 7000996 A.U. (N=20); in *msps-RNAi* stage 6-7 nurse cells, 49708953± 5241944 A.U. (N=21). An unpaired t-test with Welch’s correction was performed between control and *msps-RNAi* nurse cells, p= 0.0355 (*). (E-E’) Phalloidin and Orb staining in a Klp10A-GFP overexpressing ovariole. Oocytes are highlighted with yellow arrowheads. Scale bars, 50 µm. (F-G) Representative images of microtubule staining in control and Klp10A overexpressing egg chambers. Klp10A overexpression decreases microtubule staining in both the oocyte and nurse cells. Scale bars, 25 µm. (H) Summary of oocyte growth phenotypes in the listed genetic background (all with one copy of *maternal αtub-Gal4^[V37]^*). The numbers of control and *msps-RNAi* are the same data set used in Figure 1F. For Klp10A-GFP overexpression samples: normal oocyte growth, 0%; delayed oocyte growth, 12.1%; partial small oocytes, 12.1%; complete small oocytes, 75.8% (N=91). (I) Summary of the Orb staining phenotypes in stage 5-6, stage 7-8, and stage 9 egg chambers in listed genotypes (all with one copy of *maternal αtub-Gal4^[V37]^*). The numbers of control and *msps-RNAi* are the same data set used in Figure 1G. For Klp10A-GFP overexpression samples: stage 5-6, 92.6% Orb concentrated and 7.4% Orb dispersed (N=94); stage 7-8, 32.1% Orb concentrated and 67.9% Orb dispersed (N=84); stage 9, 16.0% Orb concentrated and 84.0% Orb dispersed (N=50).

**Supplementary Figure 3.**
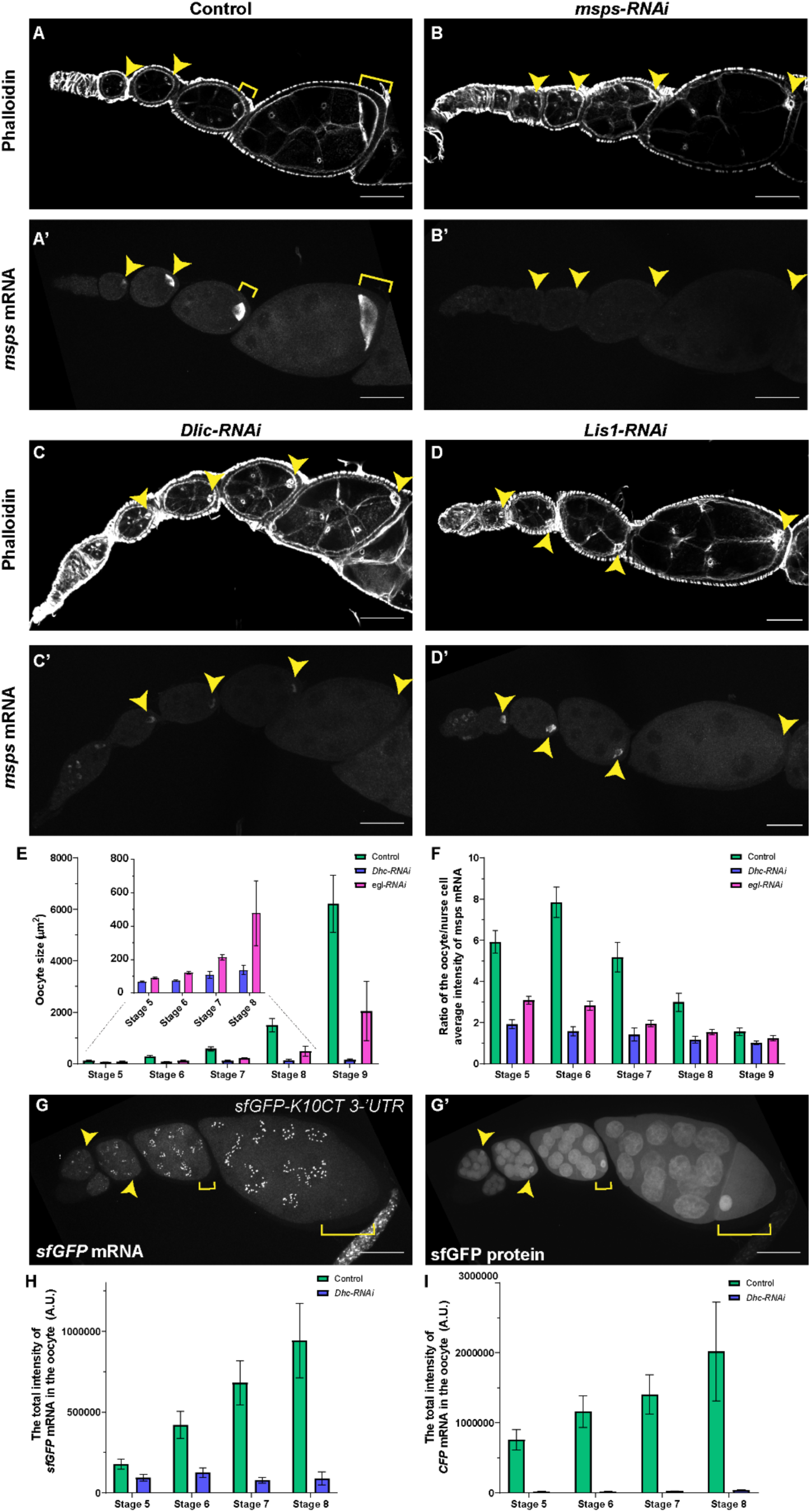
*msps* mRNA concentration in the oocyte is dependent on the function of dynein. (A-D’) Representative images of phalloidin staining and *msps* mRNA localization (by smiFISH probes against *msps* 3’UTR) in control (A-A’), *msps-RNAi* (B-B’), *Dlic-RNAi* (C-C’) and *Lis1-RNAi* (D-D’). (E) Quantification of oocyte size in control, *Dhc-RNAi*, and *egl-RNAi*. Control oocyte size: stage 5, 122.3 ± 12.2 µm^2^ (N=38); stage 6, 286.4 ± 32.8 µm^2^ (N=41); stage 7, 581.9 ± 84.7 µm^2^ (N=19); stage 8, 1496.5 ± 259.0 µm^2^ (N=23); stage 9, 6220.3 ± 1113.0 µm^2^ (N=24); *Dhc-RNAi* oocyte size: stage 5, 66.8 ± 3.9 µm^2^ (N=54); stage 6, 73.3 ± 4.9 µm^2^ (N=47); stage 7, 108.4 ± 19.9 µm^2^ (N=38); stage 8, 137.6 ± 26.7 µm^2^ (N=32); stage 9, 153.6 ± 35.9 µm^2^ (N=30); *egl-RNAi* oocyte size: stage 5, 89.2 ± 5.7 µm^2^ (N=71); stage 6, 120.8 ± 7.5 µm^2^ (N=56); stage 7, 214.0 ± 17.5 µm^2^ (N=55); stage 8, 477.7 ± 193.8 µm^2^ (N=32); stage 9, 2046.9 ± 1156.7 µm^2^ (N=25). Unpaired t-tests with Welch’s correction were performed in the following groups: between control and *Dhc-RNAi*, stage 5-stage9, all p <0.0001 (****); between control and *egl-RNAi*, stage 5-stage 9, all p <0.0001 (****); between *Dhc-RNAi* and *egl-RNAi*: stage 5-7, all p <0.0001 (****); stage 8, p=0.0012 (**); stage 9, p= 0.0025 (**). (F) Ratio of *msps* mRNA average intensity in the oocyte to nurse cells in control, *Dhc-RNAi*, and *egl-RNAi*. *msps* mRNA ratio in control: stage 5, 5.93 ± 0.54 (N=38); stage 6, 7.84 ± 0.75 (N=41); stage 7, 5.18 ± 0.72 (N=19); stage 8, 2.99 ± 0.44 (N=23); stage 9, 1.55 ± 0.19 (N=24); *msps* mRNA ratio in *Dhc-RNAi*: stage 5, 1.92 ± 0.21 (N=54); stage 6, 1.58 ± 0.22 (N=47); stage 7, 1.42 ± 0.31 (N=38); stage 8, 1.17 ± 0.16 (N=32); stage 9, 1.02 ± 0.08 (N=30); *msps* mRNA ratio in *egl-RNAi*: stage 5, 3.09 ± 0.20 (N=71); stage 6, 2.82 ± 0.23 (N=56); stage 7, 1.95 ± 0.16 (N=55); stage 8, 1.54 ± 0.13 (N=32); stage 9, 1.24 ± 0.12 (N=25). Unpaired t-tests with Welch’s correction were performed in the following groups: between control and *Dhc-RNAi*, stage 5-stage 9, all p <0.0001 (****); between control and *egl-RNAi*, stage 5-stage 8, all p <0.0001 (****), stage 9, p=0.0063 (**); between *Dhc-RNAi* and *egl-RNAi*: stage 5, p <0.0001 (****); stage 6, p <0.0001 (****); stage 7, p = 0.0031 (**); stage 8, p= 0.0004 (***); stage 9, p= 0.0034 (**). (G-G’) *sfGFP* mRNA (by smiFISH probes against the sfGFP) and sfGFP protein expressed with the C terminus of K10 3’UTR from the standard transformation pUASp vector, without the K10 TLS signal for oocyte localization (H) Quantification of the total fluorescence intensity of *sfGFP-msps 3’UTR* mRNA in control and *Dhc-RNAi*: control stage 5, 176461 ± 30966 A.U. (N=40); control stage 6, 421089 ± 83487 A.U. (N=41); control stage 7, 681176 ± 135774 A.U. (N=32); control stage 8, 942476 ± 231095 A.U. (N=10); *Dhc-RNAi* stage 5, 92832 ± 19576 A.U. (N=40); *Dhc-RNAi* stage 6, 125101 ± 29537 A.U. (N=29); *Dhc-RNAi* stage 7, 77605 ± 17385 A.U. (N=30); *Dhc-RNAi* stage 8, 89263 ± 41115 A.U. (N=15). Unpaired t-tests with Welch’s correction were performed in the following groups: between control and *Dhc-RNAi*, stage 5-stage 8, all p<0.0001 (****). (I) Quantification of the total fluorescence intensity of *CFP-Msps.CDS* mRNA in control and *Dhc-RNAi*: control stage 5, 755268 ± 145352 A.U. (N=28); control stage 6, 1162861 ± 227367 A.U. (N=35); control stage 7, 1404235 ± 279382 A.U. (N=20); control stage 8, 2020663 ± 704920 A.U. (N=14); *Dhc-RNAi* stage 5, 19164 ± 2812 A.U. (N=36); *Dhc-RNAi* stage 6, 17637 ± 2906 A.U. (N=41); *Dhc-RNAi* stage 7, 23172 ± 4154 A.U. (N=35); *Dhc-RNAi* stage 8, 36288 ± 9336 A.U. (N=18). Unpaired t-tests with Welch’s correction were performed in the following groups: between control and *Dhc-RNAi*, stage 5-stage 8, all p<0.0001 (****). All samples with one copy of *maternal αtub-Gal4^[V37]^*. Oocytes are indicated by the yellow arrowheads or yellow brackets. Scale bars, 50 µm. Data are represented as mean ± 95% confidence intervals (E-F, H-I).

**Supplementary Figure 4.**
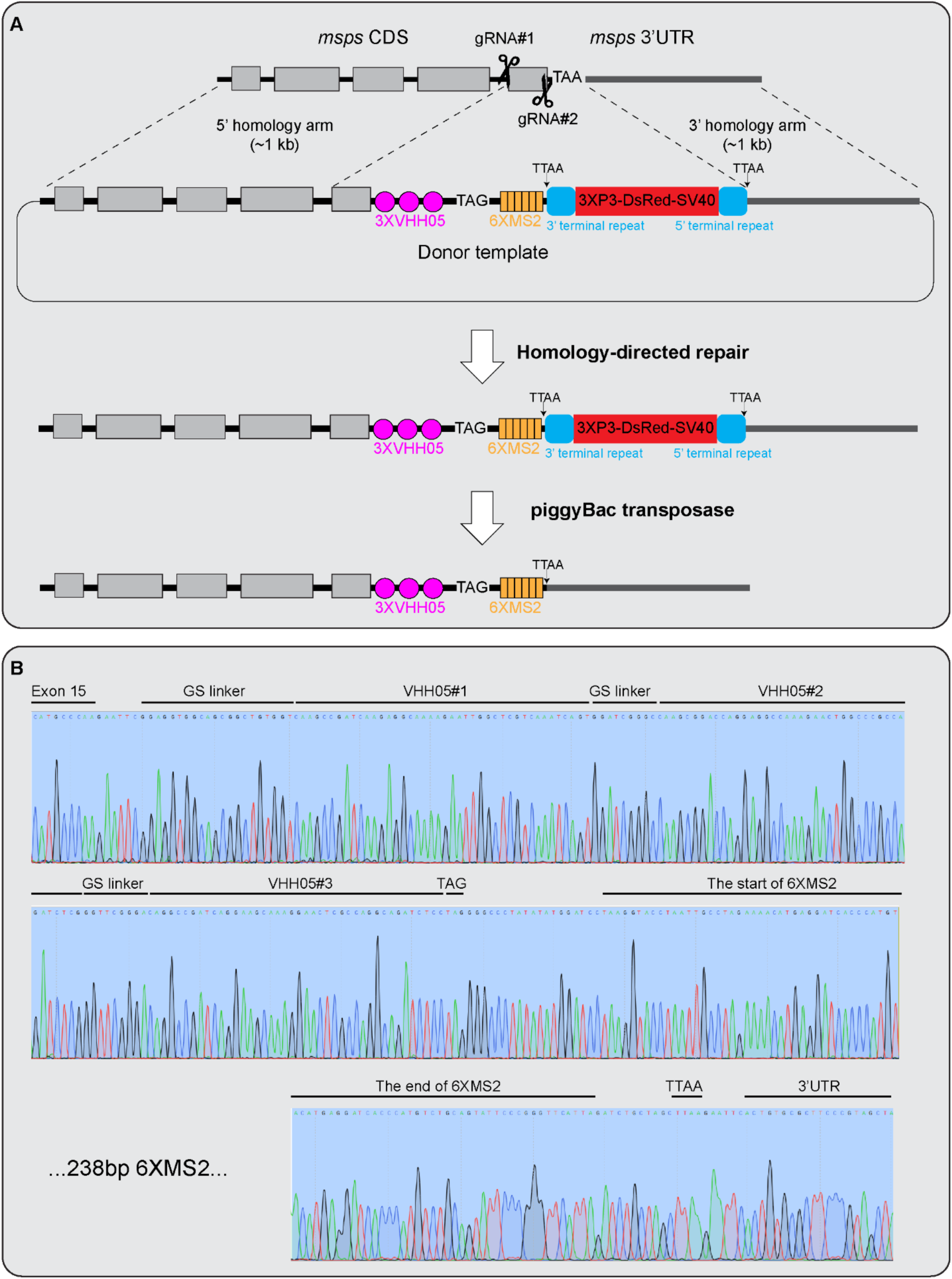
The CRISPR knock-in strategy of generating Msps protein tag. (A) A schematic cartoon illustration of the CRISPR knock-in strategy to tag the endogenous Msps protein. Two gRNAs targeting opposite strands of the last exon were used to create double-strand DNA breaks with the presence of *nos*-Cas9. A donor template containing 3 copies of VHH05, and 6 copies of MS2 sequences, followed by a fluorescent DsRed eye marker, flanked by ∼1 kb 5’ homology arm and ∼1 kb 3’ homology arm, respectively. The successful homology-directed repair was screened based on the fluorescently red eye color. The DsRed eye marker was later removed by crossing to piggyBac transposase. (B) The genomic sequences were verified by Sanger sequencing of the amplified PCR product from the extracted genomic DNA of the CRISPR knockin line of Msps-3X VHH05-6X MS2.

**Supplementary Figure 5.**
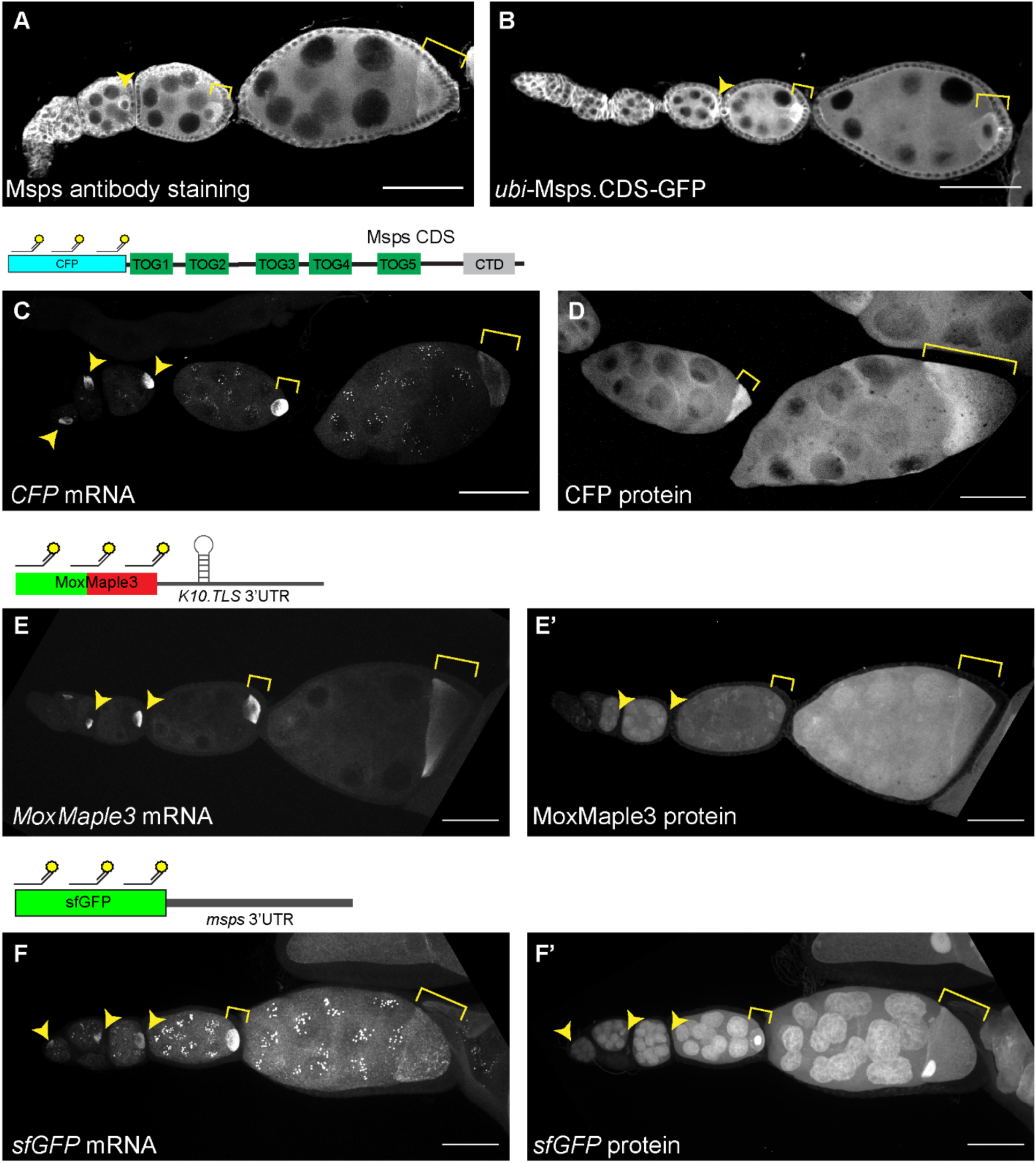
Msps protein is accumulated in the oocyte. (A-B) Msps protein is concentrated in the oocytes, shown by anti-Msps immunostaining (A) and a GFP-tagged full-length Msps under a weak ubiquitous promoter, *ubi* (B). (C-D) CFP-tagged *msps* mRNA (C) and Msps protein (D) are both concentrated in the oocytes. (E-E’) *MoxMaple3* mRNA is concentrated in the oocytes due to the addition of *K10.TLS* in the 3’UTR (E), but MoxMaple3 protein is diffused through the entire egg chambers. (F-F’) *sfGFP* mRNA is concentrated in the oocytes because of the *msps 3’-UTR* (F), but sfGFP protein is localized both in nurse cells and in the oocyte, mostly nucleus-enriched (F’). Oocytes are indicated by the yellow arrowheads or yellow brackets. Scale bars, 50 µm. (C-F’) The transgene expressions are driven by one copy of *maternal αtub-Gal4^[V37]^*.

